# Structural Insights into the Coupling Mechanism of Vectorial CO_2_ Uptake by DAB1

**DOI:** 10.64898/2026.05.01.722082

**Authors:** Naiya R. Phillips, Luke M. Oltrogge, Jonathan P. Remis, David F. Savage

**Author notes:** **Author Contributions:** N.R.P., L.M.O., and D.F.S. designed research. N.R.P. and J.P.R. prepared samples for cryo-EM and collected and processed cryo-EM datasets. N.R.P. performed molecular cloning, protein expression and purification, structural modeling and refinement, biochemical assays, and suppressor library construction and screening. N.R.P., L.M.O., J.P.R., and D.F.S. analyzed data. N.R.P., L.M.O., and D.F.S. wrote the manuscript.

## Abstract

CO_2_ transporters enable bacterial CO_2_-concentrating mechanisms (CCMs) by catalyzing directional hydration of CO_2_, yet the basis of this vectorial carbonic anhydrase (CA) activity remains an open question. We used cryo-electron microscopy to determine the structure of the DAB1 complex from *Thermocrinis albus* to 2.13 Å, revealing a heterotrimer in which a deeply buried β-CA active site in DabA is structurally coupled to the proton-translocating subunit DabB. Two conformational states define distinct solvent channels for substrate entry and product exit. A suppressor screen identifies mutations that disrupt coupling while retaining CA activity, underlying the importance of conserved residues that link proton translocation to active-site remodeling. These results support a model in which proton-driven conformational changes regulate substrate access to the active site, enabling vectorial CO₂ hydration.

**Significance Statement:** Most autotrophic bacteria rely on CO_2_-concentrating mechanisms (CCMs) to overcome the slow rate and non-specificity of the enzyme rubisco. These systems elevate cytosolic HCO_3_^-^ up to 1,000-fold higher than the environmental concentration. This steep concentration gradient requires either active HCO_3_^-^ transport or a vectorial carbonic anhydrase (CA) that couples CO₂ hydration to an energy source. The mechanism of this vectorial CA activity has remained unclear. We determine the structure of the DAB1 complex, a vectorial CA found across many bacterial phyla that is coupled to the proton motive force of the plasma membrane. We define how proton translocation remodels the buried CA active site, providing a concrete example of energy-driven catalysis and clarifying a key component of the chemoautotrophic CCM.

## Introduction

The vast majority of inorganic carbon (C_i_) enters the biosphere *via* the enzyme rubisco(1). However, rubisco is both a relatively slow and non-specific enzyme (1, 2). In addition to its carboxylation reaction, it catalyzes a non-productive oxygenation reaction (3) that then requires recycling of its 2-carbon product in a process that results in the net loss of CO_2_ (4). To overcome these impediments, many autotrophs have evolved CO_2_ concentrating mechanisms (CCMs) to saturate the active site of rubisco with CO_2_, allowing it to operate at its maximum carboxylation rate while suppressing the oxygenation reaction (5). There are multiple CCM strategies across bacteria, algae, and plants, but in general they all rely on energy coupling to transiently elevate the concentration of C_i_, which will then be converted to CO_2_ by either a decarboxylase or carbonic anhydrase (CA) in close proximity to rubisco (5, 6).

Many bacteria use carboxysomes, proteinaceous microcompartments, that colocalize rubisco and CA in their CCMs. There are two different lineages of carboxysomes: the α carboxysome encapsulates form IA rubisco and is associated with α-cyanobacteria and diverse chemoautotrophs while the β lineage contains the more plant-like form IB rubisco and is found in β-cyanobacteria. Although α- and β-carboxysomes differ in their Rubisco lineages and ecological distributions, both require one or more transporters to elevate intracellular C_i_(7, 8). In alkaline environments such as seawater, HCO_3_^-^ is the dominant species of available C_i_ and is accumulated by dedicated bicarbonate transporters such as BicA (9, 10), SbtA (11, 12), and BCT1(13), which differ in their HCO_3_^-^ affinity and energy coupling mechanisms, and are induced under different regimes of C_i_ scarcity (reviewed recently in (14)). In more acidic conditions such as freshwater or increasingly acidified oceans, CO_2_ becomes the dominant C_i_ species, making HCO₃⁻ import insufficient (15). Under these conditions, some bacteria employ transporters that directly act on CO_2_. One family of CO_2_ transporters are the specialized NDH-1 complexes found within the thylakoid membrane that bind to a soluble protein CupA or CupB (for <u>C</u>O_2_ <u>up</u>take) (16, 17). Like the different bicarbonate transporters, they are induced by different conditions, with NdhD4/NdhF4/CupB complex being expressed constitutively while the NdhD3/NdhF3/CupA complex is induced by low CO_2_ and demonstrates higher affinity. These complexes are thought to couple redox with vectorial CA activity, driving CO_2_ hydration against an existing high cytosolic HCO_3_^-^ concentration. β-cyanobacteria frequently utilize all five of these transporters, while α-cyanobacteria, depending on their environments, possess BicA, SbtA, and/or CupA (18).

Many chemoautotrophs that reside in acidic environments, and even many heterotrophs, instead use a distinct C_i_ uptake system called the DABs Accumulate Bicarbonate (DAB; also called DAC or MpsAB) complex based on its function (19–22). This complex always contains a cytosolic β-CA-like domain (DabA) and a transmembrane proton translocation subunit (DabB). In some cases, these two functional domains are fused into one long protein, though they are more often separate ORFs. Genetic assay, proteomics, and biochemistry have determined that these subunits form a requisite heterodimer with 1:1 stoichiometry (19, 22, 23). In a subset of operons that we previously termed the DAB1s, there is a small third protein called DabC that has been found to be essential in the complexes where it occurs, though its function is unknown (23). Similar to the Cups, the DABs are proposed to act directly on CO_2_ as energized vectorial CAs to capture C_i_, but rather than using redox potential, they are thought to use the proton gradient of the plasma membrane. This is supported by experiments in which cells expressing a DAB and treated with the protonophore carbonyl cyanide *m*-chlorophenyl hydrazone (CCCP) fail to accumulate C_i_, despite active C_i_ accumulation in the absence of the CCCP (22, 23).

While there have been a number of studies clarifying the mechanisms of the various HCO_3_^-^ transporter systems associated with the bacterial CCM (reviewed in (7)), the mechanism of the CO_2_ uptake systems–either the Cups or the DABs–has remained unsolved. In particular, it is an open question how these complexes mechanistically act as vectorial CAs, rather than reversibly catalyzing hydration, as all other CAs do. Recent structural and mutational studies along with MD simulations (24–27) on the Cups have granted some clues into a possible mechanism. They suggest that a novel CA active site at the interface of the NDH-1 complex and the Cup protein accepts CO_2_ entering from a hydrophobic channel through the NdhF3 subunit. The NdhF3 subunit then shuttles the H^+^ produced from the hydration reaction across the thylakoid membrane, thus disallowing the reverse dehydration. We sought to solve the structure of the DAB complex to understand how it catalyzes a similar reaction by coupling only to a proton gradient, without apparent redox involvement.

To that end, we used Cryo-electron microscopy (Cryo-EM) to solve the structure of the DAB1 complex from the thermophile, *Thermocrinis albus*, to 2.13 Å resolution. The structure suggests DabA is a novel variant of β-CA with an active site buried deep in the protein core, sequestered from bulk solvent. Focused refinement revealed two distinct conformations of the active site, which are consistent with reactant entry and product exit respectively. These data, along with additional mutational analysis, provide insight into a putative multi-step mechanism for the directional coupling of CA activity.

## Results and Discussion

### Dab1 is a heterotrimer containing DabA, DabB, and DabC

In order to determine the structure of the Dab complex, we synthesized and screened seven DAB homologs for function and expression level using an *E. coli* strain lacking CAs (BW25113 *ΔcanΔcynT*), CAFree. Without a CA, the rate of equilibration of atmospheric CO_2_ to HCO_3_^-^ is too slow to provide enough cytosolic HCO_3_^-^ flux to support central metabolism, thus rendering the strain unable to grow in atmospheric air unless expressing a functional C_i_ importer or CA. All but two homologs were able to rescue CAFree. The DAB1 operon from *T. albus* (Figure 1A) appeared to have the most homogeneous and stable expression via western blot analysis (Supplementary Figure 2) and was chosen for further structural analysis.

**Figure 1.**
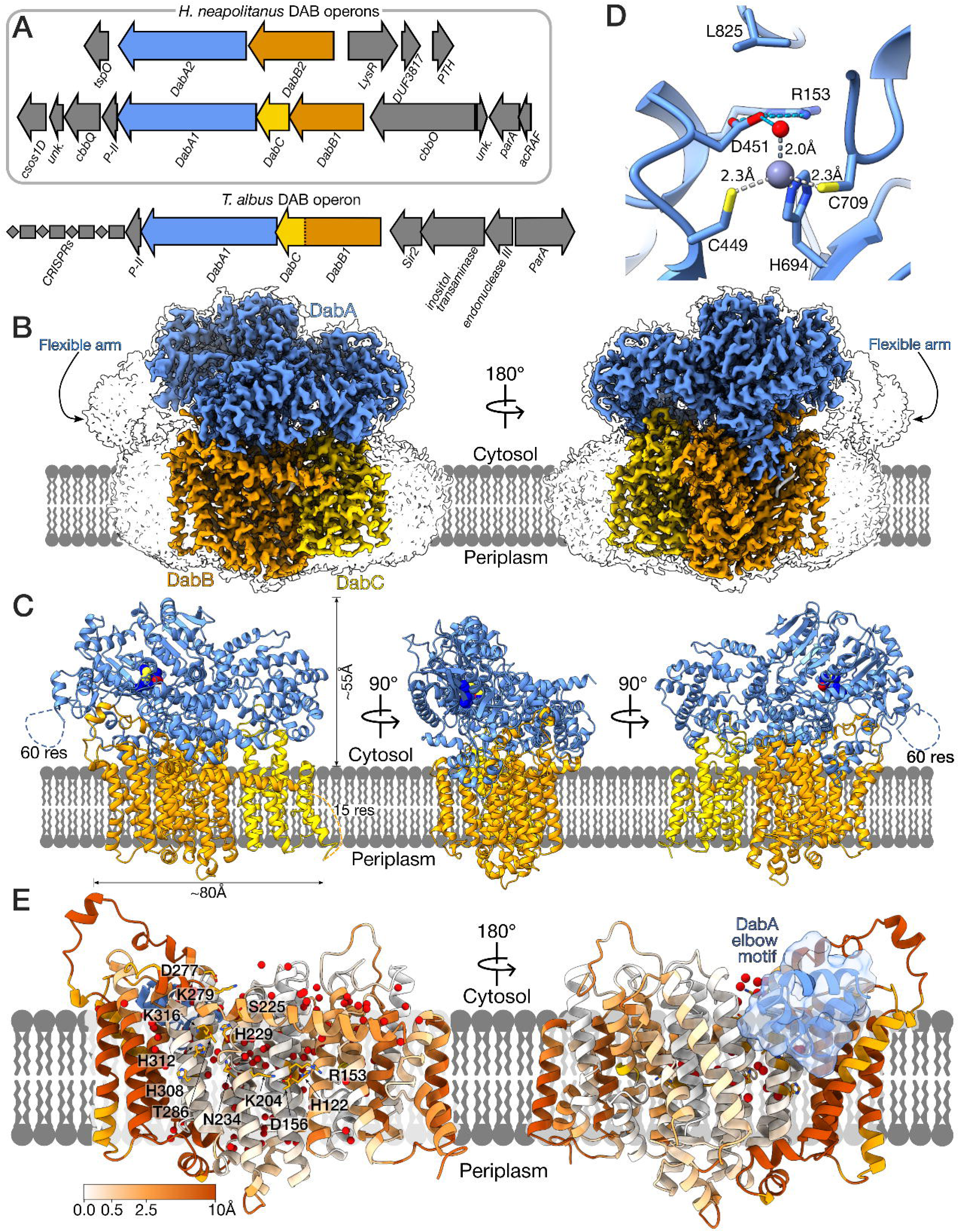
Structural features of the DAB1 Complex. (A) Comparison of the DAB1 and DAB2 operons from H. neapolitanus with the DAB1 operon from T. albus. (B) Different views of the cryo-EM density map of the T. albus DAB1 complex, segmented by subunit. The detergent micelle and flexible arm are shown as a low contour transparent surface (cryoSPARC threshold 0.0135) and the membrane is drawn schematically. (C) Different views of the atomic model of the DAB1 complex. Unmodelled residues are shown as a dotted line and active site residues are displayed as spheres. (D) A Zn^2+^ ion is coordinated by conserved β-CA active site residues and water in the DabA structure. A conserved Leu replaces a position that is typically occupied by a hydrogen bond donor that stabilizes bicarbonate in typical β-CAs. (E) Structure of DabB and DabC colored by RMSD relative to NuoL and NuoM respectively (PDB:7Z7S). Conserved proton translocation residues and waters (red spheres) are highlighted.

We expressed the DAB1 complex from *T. albus* with a C-terminal His-tag on DabC in BL21-AI *E. coli*. The complete 200kDa complex could be solubilized from the *E.coli* membrane with the detergent Lauryl Maltose Neopentyl Glycol (LMNG) (28) and purified by Ni-IMAC, heat treatment at 70°C, and gel filtration chromatography (Supplementary Figure 2), demonstrating a strong association between the transmembrane and cytosolic domains.

Cryo-EM analysis revealed a structure in which DabB and DabC are present in the detergent micelle as transmembrane proteins, as predicted by DeepTMHMM (29), while DabA is primarily cytosolic and interacts with DabB and DabC at the surface of the membrane (Fiigure 1B-C). DabA also extends two hinged helices, referred to here as the elbow motif (residues 534-565) (aka “finger-like motif” in (30)), which drops into the membrane and wedges into DabB (Figure 1E, Supplementary Figure 12). The elbow is connected to a cytosolic arm (residues 580-661) that was notable among all other regions of the complex as too flexible to model. While the majority of the map is resolved to <2.2Å (Supplementary Figure 3), the residues 590-650 of the flexible arm are visible only at a low contour (Figure 1B-C). These data are generally consistent with the predicted AlphaFold 3 (AF3) (31) model, although there are considerable differences in the shape of the elbow motif and in the region surrounding the putative active site (Supplementary Figure 5).

### DabA is a novel β-carbonic anhydrase fold

Previous work in our lab demonstrated that DabA possesses a β-CA-like Zn-containing active site that when mutated abolishes the ability to rescue CAFree (22). Our structural analysis confirms that these residues do in fact form a typical β-CA active site in which a Zn^2+^ ion is coordinated by two cysteines (C449 and C709), a histidine (H694), and a water in a tetrahedral geometry (Figure 1D). Despite sharing only ∼20% sequence homology, the closest structural homolog to DabA found by Foldseek (32) is the α-carboxysomal CA, CsosCA from *H. neapolitanus*. This suggests a possible evolutionary relationship between CsosCA and DabA—likely either DabA evolved from CsosCA, or both proteins diverged from a common ancestral β-CA. In both the structure of DAB1 and of CsosCA (PDB: 2FGY), the Zn-bound water is stabilized by a nearby aspartate (D451 in DAB1), which is itself stabilized by hydrogen bonding with a nearby arginine (R453) (33). However, the position where other β-CA active sites have another hydrophilic residue, such as glutamine (34) or histidine (33) to hydrogen bond with HCO_3_^-^, is occupied by a leucine, L825 in *T. albus* DabA, that is broadly conserved amongst DabA orthologs (Figure 1D, Supplementary Figure 6). It is possible that this lack of an H-bond donor in DabA1 raises the barrier to HCO_3_^-^ binding and encourages product release after hydration.

Unlike typical β-CAs, the DabA active site is found deep within the protein core. Most other known CAs have active sites located close to the protein surface where they are directly accessible to substrates CO_2_ and HCO_3_^-^ (reviewed in (35, 36)). In contrast, the DabA active site is buried more than 10 Å from solvent, sequestered by a flexible lid domain comprising residues 471–509 (Figure 3B). Although we observe multiple conformational states for the lid (see below) and surrounding environment, the active site always remains distant and restricted from the solvent. Specifically, CAVER 3.0 (37) analysis reveals that accessing the active site requires traversing channels 12–20 Å in length (Supplementary Figure 7A).

**Figure 2.**
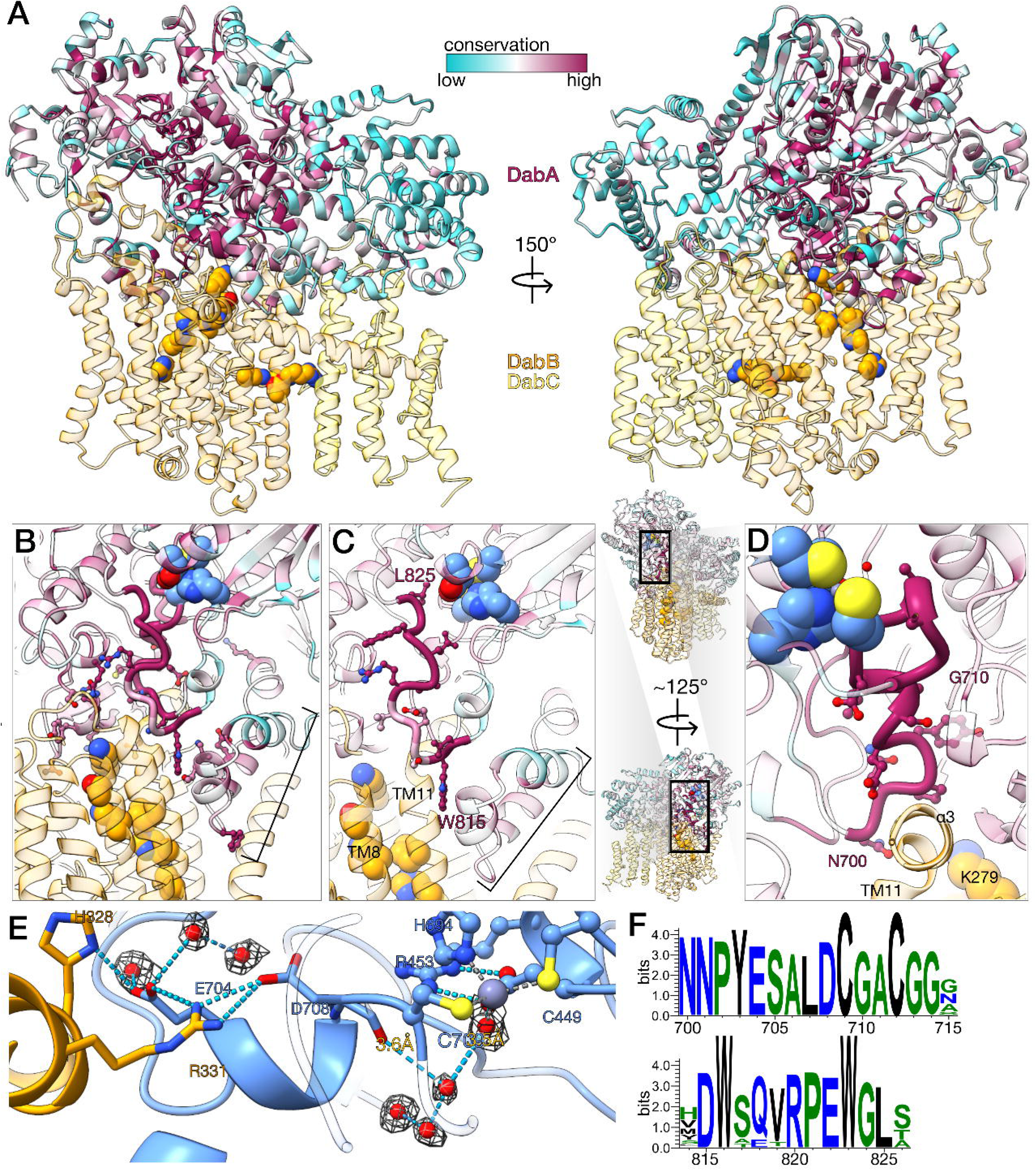
Conserved structural components of the novel DAB fold. (A) Structure of the DAB1 complex with DabA colored according to sequence conservation. (B) A view of the conserved contact residues within DabA. Contacts were calculated within ChimeraX using the criteria of ≥15Å^2^ of buried solvent accessible surface area, and contacts with greater than 80% sequence conservation were displayed. Proton translocation residues in DabB and active site residues in DabA are displayed as spheres. The elbow motif is indicated by a bracket. (C) Structure of the DW(XXX)RPEWGL motif. Residues of note are W815 contacting both DabA and B and L825 within the active site. (D) Structure of the NNPYESALDCGACGG motif, including the active site C709. (E) Hydrogen bonding interactions between the DabB α3 helix and E704 and D708 within the NNPYESALDCGACGG motif proximal to the active site. (F) WebLogo showing degree of conservation within the displayed motifs.

**Figure 3.**
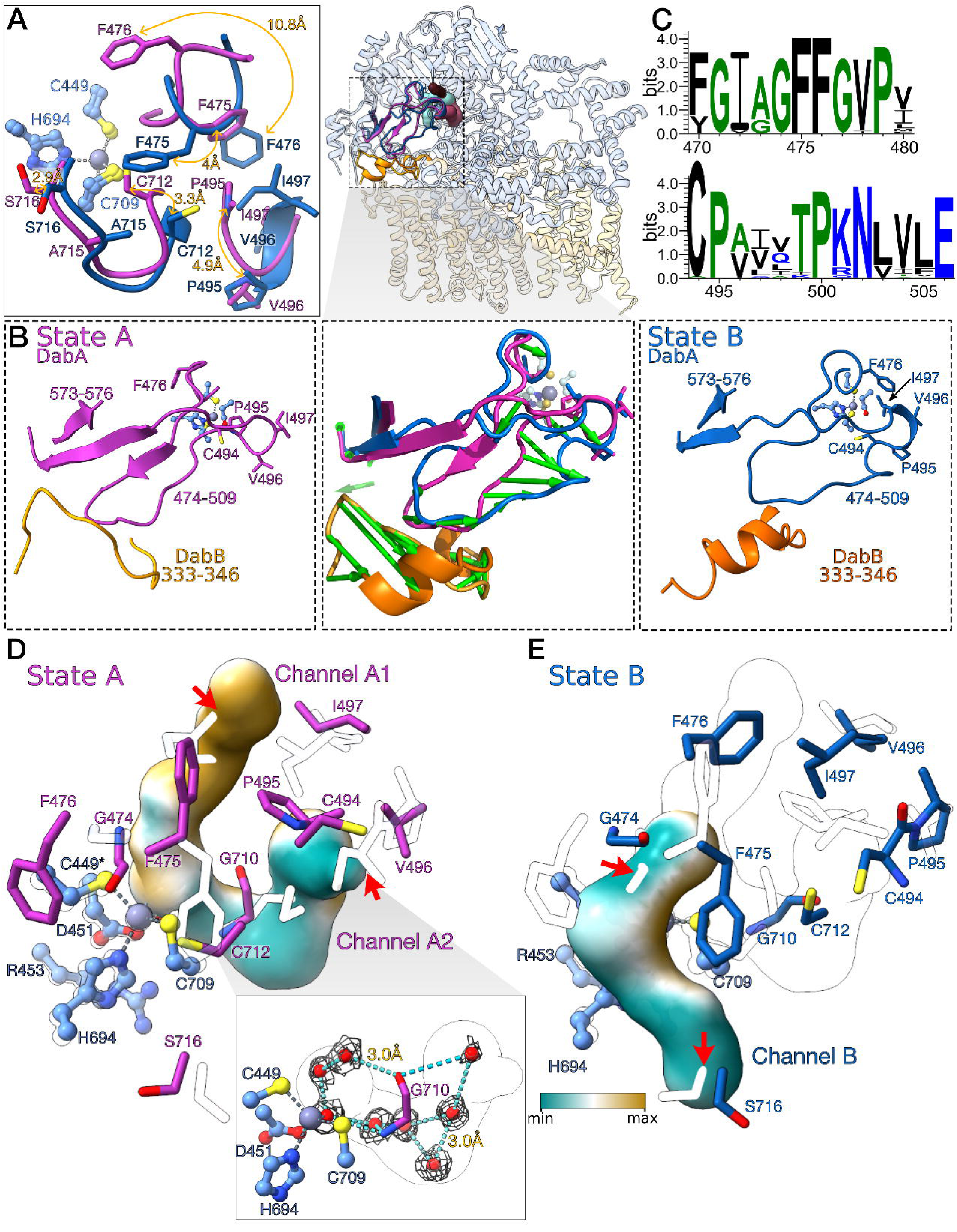
Structural comparison of State A and State B reveals distinct substrate and product channels. (A) Local structural changes surrounding the DabA active site in State A (magenta) and State B (dark blue). Distances are shown between the atoms indicated by the arrows to highlight key movements between the two states. (B) Remodeling of the lid region enclosing the active site (residues 474-509) and coordinated movement of DabB (residues 333-346). The top panel shows the location of this region within the full complex. The center panel illustrates the principal motion between State A (left) and State B (right), calculated using the modevectors PyMOL script to highlight the direction and magnitude of the residue displacements. (C) Sequence conservation of mobile residues within the lid, shown as WebLogos. (D) State A contains two channels, colored by the hydrophobicity of the surrounding protein surface. Channel A1 is hydrophobic while Channel A2 is hydrophilic. The State B conformation is shown semi-transparent to reveal how the residues are positioned in State B to close Channel A1 and A2. Red arrows highlight the role of F476 in closing Channel A1 and of C494 in blocking Channel A2. Inset shows a chain of ordered waters and their EM density lining Channel A2. (E) State B contains a distinct hydrophilic channel that opens to the opposite face of DabA as channels A1 and A2. This channel is blocked in State A by G474 and S716, highlighted by red arrows.

In addition to the five active site residues conserved in both DABs and CAs, there are three highly conserved (see Methods for additional details) DAB-specific motifs near the active site (Figure 2A). First, the 471-GI(A/G)GFFG-477 motif within the lid domain forms a mobile loop to apparently modulate active site accessibility. A second longer motif, 700-NNPYESALDCGACGG-714, is strictly conserved across 50 diverse DabA1 sequences, with the motif containing substitutions in the YESA residues in DabA2 sequences. This loop extends from the top of the transmembrane 11 (TM11) helix of DabB (see Supplementary Figure 9 for helix numbering) all the way to the active site (Figure 2D). Side chains and backbone atoms within this loop form hydrogen bonds with both DabB and structured waters in the active site. Similarly, the 815-DW(XXX)RPEWGL-825 motif inserts W816 between the DabA elbow motif and DabB TM11 (Figure 2C). It then spans the distance to the CA active site where L825 is positioned to prevent stable HCO_3_^-^ binding as discussed above. The structures of these two longer motifs connecting DabA and DabB are well conserved between *T. albus* DAB1 and *H. neapolitanus* DAB2 (Supplementary Figure 13E).

### DabB demonstrates homology to *E.coli* NuoL

DabB is homologous to the NADH:quinone oxidoreductase/Mrp antiporter protein family (PFAM: PF00361) and displays strong structural homology to the NuoL subunit of E. coli complex I (PDB:7Z7S) (RMSD between 277 pruned atom pairs is 1.18 Å) (Figure 1E). NuoL is thought to primarily contribute to proton transport via a network of water molecules and hydrophilic residues embedded in the interior of the membrane (38, 39). DabB likely also allows for proton transport *via* a similar mechanism. The residues involved in proton transport are conserved (Supplementary Figure 9) and interact with structured waters that form a chain through the membrane, and the helices containing these residues are highly structurally conserved between DabB and *E. coli* Complex I (39). The helices that do not contain any of the conserved proton translocation residues are disrupted by DabA’s membrane helices.

### DabB and DabA swap helices to transmit conformational changes

DabA interacts with DabB through extensive contacts along the cytosolic face of DabB—many between conserved residues (Figure 2A-B)—and through the elbow motif (Supplementary Figure 12). This motif inserts ∼20Å into the membrane and displaces TMs 12-15 relative to their position in NuoL. Lo *et al.* propose that this insertion allows the elbow to participate in proton translocation (30). In our model, however, none of the residues within this motif approach the proton-translocation pathway closely enough to play such a role (Supplementary Figure 13C). The elbow does interact with the TM11 helix that contains the highly conserved proton translocation residues H308, H312, and K316 (Figure 2B-C), although it does so without disrupting structural homology to NuoL; the structure of TM11 remains highly conserved within the transmembrane region (RMSD < 0.5Å between DabB and NuoL) (Figure 1E). In turn, the TM11 helix then extends above the membrane and is attached by loops to short cytosolic helices that are absent in NuoL. These helices interact closely with a highly conserved motif in DabA: D708 and E704 within the DabA motif hydrogen bond with conserved residues R331 and H328 in DabB as well as several ordered waters (Figure 2E). As this interaction is proximal to the active site C709, these interactions likely allow DabB to directly influence CA activity.

### DabC is likely a structural rather than catalytic subunit

DabC is a distinguishing characteristic of DAB1 operons relative to DAB2 operons. Unlike DabB, DabC does not belong to any previously described PFAM domains. Although it is only about half the size of NuoM, its helices do structurally align with the NuoM helices proximal to NuoL in Complex I (39). While NuoM contributes to proton translocation in Complex I *via* conserved hydrophilic residues embedded in the transmembrane helices, those residues were found to be hydrophobic in DabC and we did not observe the characteristic chain of structured waters through the protein, suggesting that it does not contribute to proton transport. However, a CAFree assay of *ΔdabC* mutants of both *H. neapolitanus* and *T. albus* indicates the protein is required for function *in vivo*. All together, this suggests that DabC may play a structural or allosteric role in the DabABC complex, rather than a catalytic one (Supplementary Figure 10).

### Two conformations of the active site regions can be resolved to high resolution

The active-site region was unexpectedly poorly resolved despite the high overall map quality, suggesting underlying structural heterogeneity. We therefore performed 3D classification, which revealed two distinct states with sufficient quality cryo-EM density for model building. In these conformations, the active site of Zn^2+^ and its coordinating residues show little variation while surrounding residues move dramatically, leading to two different states of solvent accessibility to the active site (Figure 3A-E, Supplementary movie 1). These two states likely represent two of several states the complex undergoes in a complete catalytic cycle.

The first model, referred to as State A (164,827 particles), bears a closer resemblance to the predicted AF3 structure (Supplementary Figure 5). The highly conserved GI(A/G)GFFG motif (Figure 3B-C) is in a coiled position and the remaining lid residues (480–509), adopt a structured β-strand conformation with the highly conserved C494 and P495 residues present in a loop between two strands. CAVER analysis of this conformation revealed two putative substrate channels between the active site and the cytosol (Figure 3D). Channel A1 is formed primarily from conserved hydrophobic side chains and terminates at the active site Zn^2+^, where a coordinated water is observed. We hypothesize this is a putative CO_2_ channel, as its path would place any entering CO_2_ molecules in position for attack from the hydroxide ion formed during catalysis. We were unable to resolve any distinct density that could be unambiguously assigned as CO_2_ (Supplementary Figure 7), however Lo *et. al* did capture putative CO_2_ molecules in the equivalent channel in DAB2 (30)(Supplmentary Figure 13B). Channel A2 is more hydrophilic, and we observe a chain of distinct structured densities, which we interpret as water molecules, suggesting the channel functions to replenish the water used in catalysis (Figure 3D inset). Overall, we hypothesize that the State A conformation could allow entry of both reactants to the active site.

In the second state, State B (168,739 particles), the β-strands capping the active site transition to an irregular coil structure which remodels the active site channels (Figure 3B,E). Relative to State A, the GI(A/G)GFFG motif adopts a more helical conformation, forcing F476 to undergo a dramatic movement (Cα change of ∼5Å, Cγ change of ∼11Å) to close off the hydrophobic Channel A1, and C494 moves ∼5Å between to block off the water entry Channel A2 with its Cα and backbone nitrogen. Meanwhile, both G474 and S716 shift ∼3Å to open a new hydrophilic channel (Channel B) opposite to the State A channels, which opens to the solvent near the membrane leaflet. We propose that this channel could represent an HCO_3_^-^ exit channel following catalysis (Fig 3C). Together, these structures support a model in which the substrates enter through CO_2_ and water channels for CO_2_ hydration, after which a transition to State B closes the entry routes and opens a separate exit channel to release HCO_3_^-^ to the cytosol.

### A library screen reveals two key features of coupled activity

While our structural data and previous functional data are consistent with vectorial CA activity, the mechanism coupling DabA and DabB functions remains unknown. We investigated the molecular basis of this mechanism using a suppressor screen in CAFree (Figure 4A, Supplementary Figure 15A). Briefly, we mutated the essential proton-translocation residue H229 in the *T. albus* DabBC protein (equivalent to H260 in *H. neapolitanus* DabB2), which abolished proton conduction and rendered either complex unable to rescue CAFree growth in air (Figure 4B). This complete loss of activity indicates that DabA catalysis is strictly coupled to proton translocation. We then generated error-prone PCR libraries of *H. neapolitanus* DabA2 (∼1.2×10^6^ variants) to express with *Hn*DabB2-H260A and of *T. albus* DabA1 (∼1.4×10^6^) to express alongside *Ta*DabBC-H229A and screened for variants that rescued growth in ambient air. We expected that these variants would have broken the strict coupling between DabA and DabB, allowing DabA to act as a non-vectorial CA that rescues CAFree growth, and that the mutations enabling this rescue would identify residues critical for the coupling mechanism.

**Figure 4.**
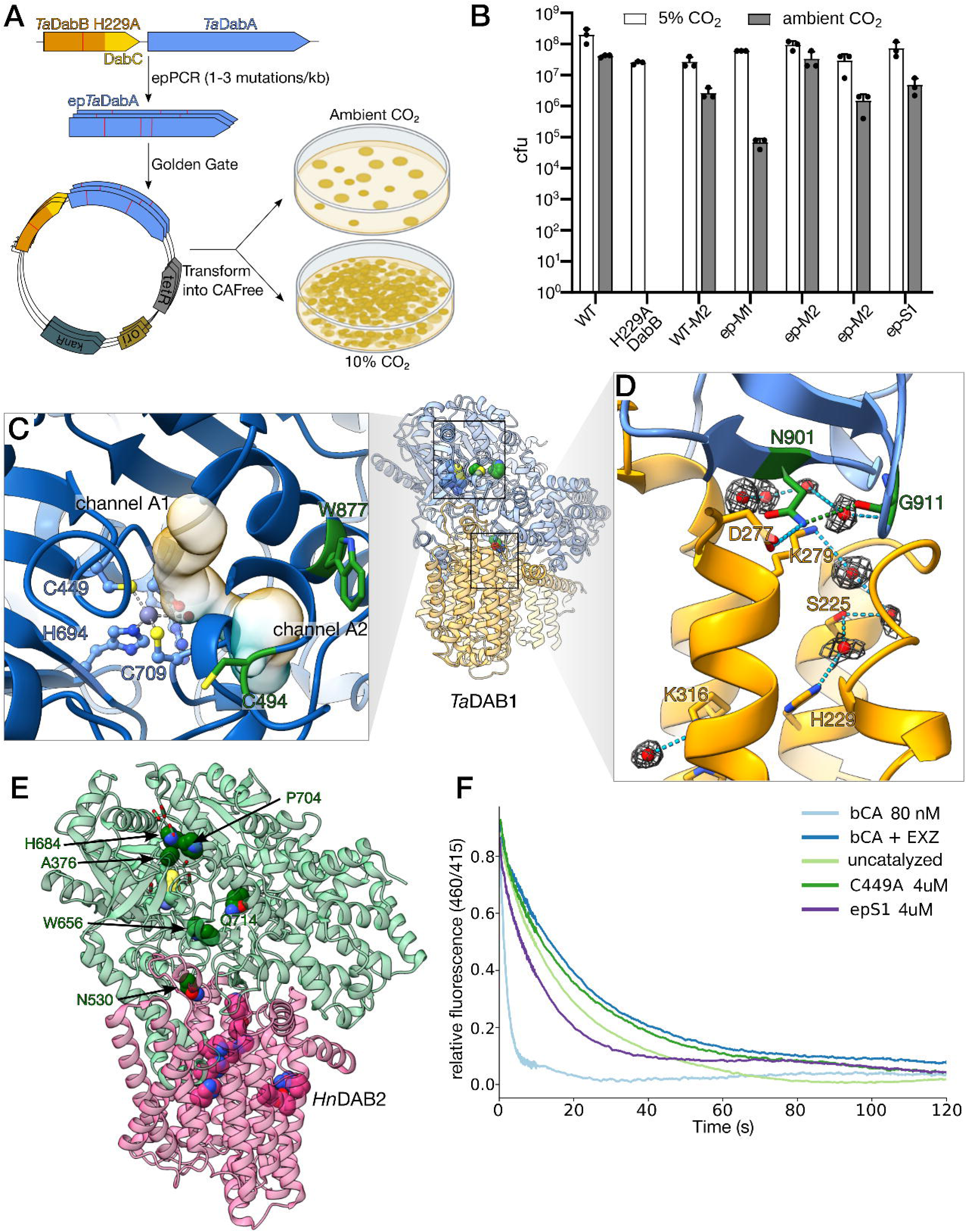
A suppressor screen highlights two key structural features for coupling DabA and DabB. (A) Schematic of the suppressor screen used to identify mutants in which DabA CA activity is decoupled from DabB proton translocation. (B) Growth of CAFree expressing decoupled *T. albus* variants in 5% CO_2_ (permissive) and ambient air (selective). (C) Structure of the *T. albus* active site shown in State 2 with residues identified in the screen highlighted in green. C494 and W877 exemplify mode 1 of decoupling: preventing proper channel closure during the catalytic cycle. (D) Structural view of the DabA:DabB interface with residues identified in the screen highlighted in green and conserved residues in the DabB proton translocation pathway shown. These mutants illustrate mode 2 of decoupling: disrupting signal transmission from DabB to DabA during proton translocation. (E) Residues from the *H. neapolitanus* DAB2 screen mapped onto the model of the DAB2 complex (PDB:9RD0). Each substitution is individually sufficient to decouple DAB2 activity, and they are displayed together for clarity. As in *T. albus*, mutations cluster near the predicted active site or between it and the proton translocation pathway. (F) A fluorescence-based carbonic anhydrase assay of the epS1 variant shows that it increases the rate of CO_2_ hydration relative to the uncatalyzed reaction, but remains substantially slower than bovine CA (80 nM). The active site mutant C449 and bCA inhibited by ethoxyzolamide (EXZ) are included as negative controls. Curves are smoothed averages of 4 time courses, with background fluorescence subtracted.

The *H. neapolitanus* error-prone library produced ∼200 colonies that grew in air, while the *T. albus* produced 5 (Figure 4B). Alongside the epPCR, we amplified the DabA genes with a high-fidelity polymerase to screen separately to acquire an estimate of background mutation rates. In both cases, a small number of colonies also grew on these background plates. All *T. albus* and thirty *H. neapolitanus* colonies were subjected to plasmid sequencing. Plasmids containing any mutations to DabB were excluded from further analysis, giving the final list of unique sequenced variants presented in Table 1 and Supplementary Table 2.

**Table 1.** *T. albus* DabA variants with uncoupled CA activity.

| Variant | Mutations present |
| --- | --- |
| WTM2* | R66S, C494F |
| epM1 | R66S, S521C, I610F, R648G, N901K, G911D |
| epM2 | R66S, F670L, F724I, N901H |
| epM3 | R66S, R144H, H273Q, D388N, A520V, W877S |
| epS1 | D45N, R66S, C494W, T760S, A789T |

\*Colonies are named based on whether they originated from the high fidelity (WT) or error-prone (ep) amplification, and their relative size (B for big, M for medium, S for small).

Strikingly, many *H. neapolitanus* variants contained only one non-synonymous mutation to DabA, whereas every *T. albus* variant contained multiple mutations. These mutations were not isolated to any one region of the protein, suggesting that coupling is enforced by a network of rigid structural interactions between DabA and DabB throughout the complex. This model of coupling would also explain why multiple mutations are required for decoupling the complex from the thermophile *T. albus:* its inherent thermostability likely preserves the integrity of the interaction network when screened at 37°C, making single substitutions insufficient to disrupt coupling. There are likely many ways of interrupting this network that would result in a catalytically dead complex. For example, if DabB cannot force DabA to shift conformations, it could become trapped in one conformation, unable to complete a catalytic cycle. Our screen reveals substitutions that instead relax the constraints normally imposed by DabB while preserving catalytic activity in DabA. These mutants highlight two features of the complex that are required to normally maintain coupling: regulation of channel opening and closure, and mechanical linkage to transmit structural changes from the proton translocation pathway to the DabA active site.

#### Feature 1: Controlling channel closure

The importance of properly closing substrate channels during catalysis is most clearly demonstrated by the WTM2 variant, as it only contains C494F and R66S mutations. C494 is highly conserved in DabA1s and its backbone shifts ∼4Å between States A and B to close the putative water channel to the active site. Substitution to a bulky aromatic sidechain could reduce backbone mobility and interfere with channel closure, leading to unregulated solvent access (Figure 4C). Accompanying the C494F mutation is R66S, which was identified in all the *T. albus* variants. R66 is not a conserved position and is distal to both the active site and the DabB interface, but it was present in a low abundance in the initial plasmid stock used for epPCR. Its repeated selection in our screen suggests it modestly perturbs the stability of the complex, making the coupling network more susceptible to disruption by additional mutations.

Two other variants also highlight the importance of channel closure: epS1, which contains a C494W mutation, and epM3, where a W877S mutation on the surface near the hydrophilic channel exit may keep the channel constantly open. Further evidence comes from residues found in the *H. neapolitanus* screen considered in the context of the recently published structure of the *Hn*DAB2 complex (30). A376, P704, and H684 are all residues lining the CO_2_ channel near the surface, and are all found within <4Å of a CO_2_ molecule within that channel (Supplementary Figure 14C). Furthermore, A376 is part of the GI(A/G)GFFG motif that moves to seal the CO_2_ channel in State B. Any of these mutations could prevent proper closure of the CO_2_ channel during the catalytic cycle.

#### Feature 2: Mechanical linkage of proton translocation and the CA active site

A second feature of the structure highlighted by the screen is the mechanical link between the DabB proton translocation pathway and the DabA active site. This link is likely defined by a set of conserved residues at the DabA-DabB interface near the top of the proton translocation pathway, as well as the two conserved motifs that connect these interfaces to the DabA active site (Figure 2B-D). This importance of this linkage is exemplified by the epM1 and epM2 variants, which contain N901K and N901H mutations respectively. N901 is a conserved residue that bridges the two subunits by hydrogen bonding with the conserved proton translocation residue D277 in DabB, and with the backbone carbonyl of G911 in DabA, which is mutated to G911D in epM1 (Figure 4D). Altering these residues could effectively disconnect DabA from proton translocation in DabB. Mutations from the *H. neapolitanus* screen that reinforce the importance of these links are N530D (corresponding to N700 in *T. albus*) and W656R/C (W823) (Figure 4E). Both of these residues are strictly conserved in the sequence motifs that span between DabB and the active site, and their mutation likely impairs their ability to propagate structural changes between the two proteins.

### *In vitro* CA activity assays demonstrate modest uncoupled activity

We sought to confirm that these DabA variants function as non-vectorial CAs using stopped-flow kinetic assays. Hydration kinetics were measured by mixing the enzyme with a saturated CO_2_ solution and monitoring the resulting pH-dependent fluorescence shift (40). Only one tested variant, epS1, which contains a mutation to C494 that likely perturbs the State A - State B transition, demonstrated *in vitro* CA activity. Although we only observed a modest improvement in *v*_0_ over the uncatalyzed reaction (Figure 4F), it is the first confirmation of *in vitro* activity in a DAB and supports the hypothesis that vectorial activity can be uncoupled by mutation (22, 41). We suspect that the apparent low catalytic rate likely results from the complex’s instability outside the membrane environment.

### A model for DAB1 function

The presence of at least two distinct structural states with different solvent access channels, together with our mutational data underscoring the importance of properly closing these channels, is suggestive of a model for the hypothesized vectorial CA activity of the DAB1 complex (Figure 5).

**Figure 5.**
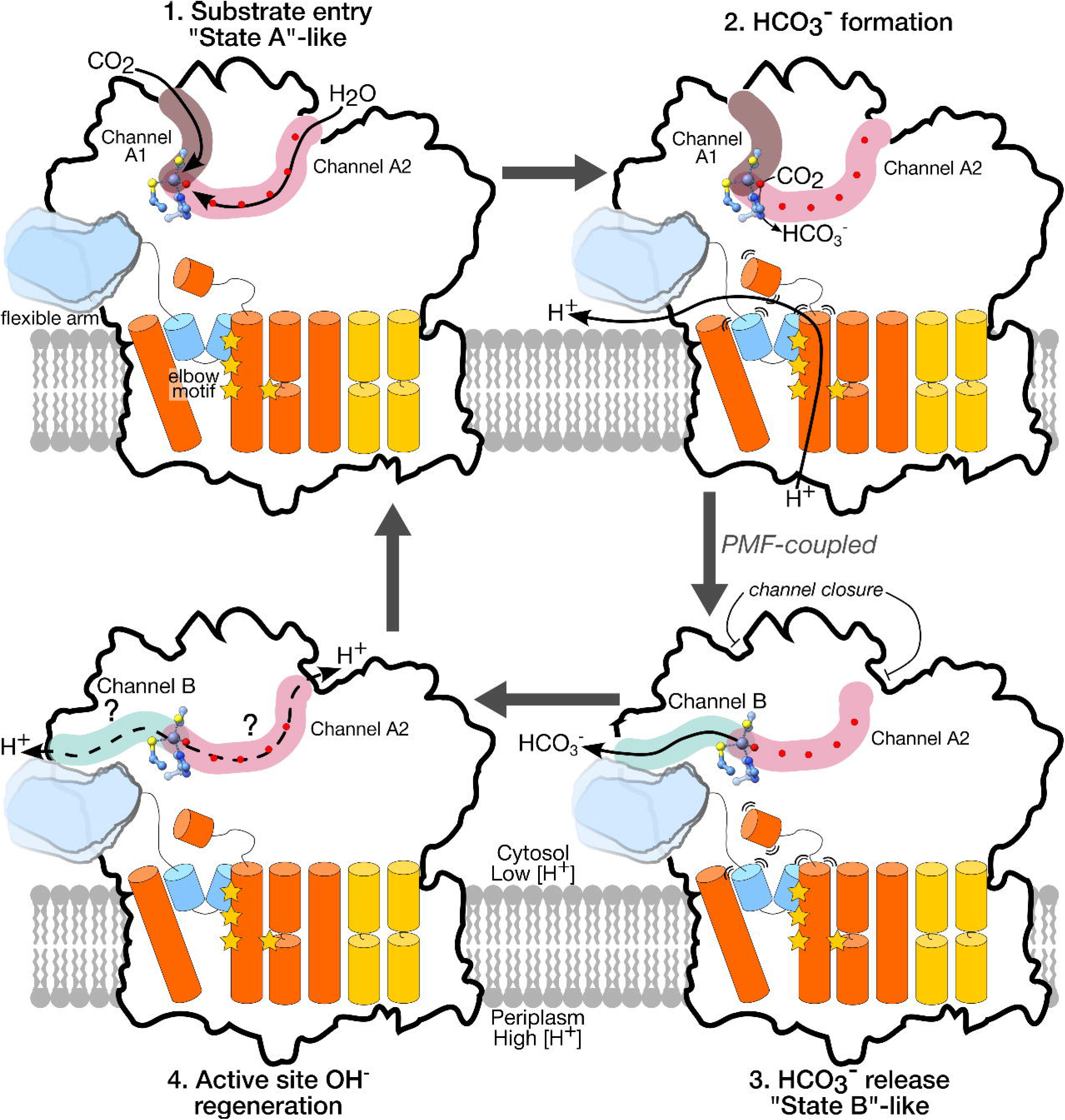
A model for DabA vectorial carbonic anhydrase activity. In State A, two channels open to the cytosol, allowing for substrate entry and catalysis of the hydration reaction. Proton translocation through DabB, mediated by conserved residues (yellow stars), triggers a conformational shift to State B, in which a distinct channel opens to allow for HCO_3_^-^ release. A new water molecule must bind the active site Zn^2+^ ion and be deprotonated to reset the complex for another catalytic cycle. Possible routes of H^+^ removal are shown as dashed lines.

The catalytic cycle for CO_2_ hydration in β-CAs can be described in four steps: 1) CO_2_ entry, 2) nucleophilic attack by the Zn^2+^-bound hydroxide to form HCO_3_^-^, 3) HCO_3_^-^ release and H_2_O rebinding, and 4) deprotonation of Zn^2+^-bound water to regenerate the hydroxide (35). The conformational features of State A are consistent with the first two steps. The CO_2_ and water channels are open and the Zn^2+^-bound water is positioned for nucleophilic attack by D451. In typical CAs, HCO_3_^-^ diffuses to bulk solvent immediately after its formation, but the deeply buried active site of DabA likely precludes such immediate release. In State A, the hydrophobic CO_2_ channel offers no favorable interaction for HCO_3_^-^ and the hydrophilic channel is occluded by a structured water network (Figure 3D inset). Thus we propose that following HCO_3_^-^ formation, DabB induces the complex to shift toward a conformation resembling State B, in which the substrate entry channels close and a new hydrophilic channel opens to provide a route for HCO₃⁻ export.

Our structural data do not provide an obvious mechanism for the final step in the cycle. In the β-CA of *Arabidopsis thaliana*, deprotonation is mediated by H216 and Y212, which shuttle the H^+^ to bulk solvent through an ordered water network (42). A similar role has recently been proposed for H397 in CsosCA (43), the closest structural homolog to the DabA, but the equivalent position is occupied by L825, suggesting DabA must use a different strategy for proton removal.

Several possibilities are consistent with our structure. In State A, the chain of ordered waters extending from the Zn^2+^-bound water to bulk solvent through Channel A2 (Figure 3D inset) could support proton transfer without a dedicated shuttling residue through a Grothuss mechanism (44). Alternatively, the water stabilized by the carbonyl of G710 is positioned to accept the H^+^ and then diffuse out through the same hydrophilic channel as HCO_3_^-^, though it is important to note that in this scenario, the release of HCO_3_^-^ and H^+^ would have to occur as separate events, or else the two species could spontaneously convert back to CO_2_ and H_2_O. A third possibility is that proton transfer is mediated by D708, a highly conserved residue that bridges DabA and DabB: its side-chain carbonyl interacts with the α3 helix of DabB, while its backbone carbonyl binds a water adjacent to the Zn^2+^-bound water (Figure 2E). Such an arrangement could allow DabB direct control over whether DabA regenerates its active site. Once the catalytic cycle is complete, the complex will have captured one molecule of HCO_3_^-^ at the cost of depleting the cell’s PMF twice: once for the H^+^ that is admitted by DabB and again for the H^+^ that is generated in the cytosol by the hydration reaction. The cell must then regulate the activity of its respiratory complexes to regenerate the PMF and maintain a cytosolic pH that supports a high concentration of HCO_3_^-^.

Regardless of the mechanism of proton transfer, the complex must ultimately relax back from its product release state to a conformation resembling State A. We propose that this reset disfavors HCO₃⁻ re-entry and the reverse dehydration reaction: the HCO₃⁻ channel is closed, the hydrophobic CO₂ channel is unlikely to admit any ionic species, and the water channel maintains a structured chain of ordered waters that would exclude HCO₃⁻.

The structural context of the elbow motif and its attached flexible arm suggests that they have a role in coupling, although our data cannot resolve what that role is; the elbow is static between State A and State B, and the arm is too mobile to be visualized. Even so, the elbow penetrates ∼20 Å into DabB and lies beside the conserved proton-pathway helices, while residues 573-576 within the loop connecting the elbow to the arm form a short β-strand that contributes the small β-sheet observed in the lid domain in the State A conformation (Figure 3B). This arrangement positions both elements where they can participate in the coordinated conformational changes we propose control the catalytic cycle.

Taken together, this model demonstrates that the DAB1 complex achieves vectorial CA activity through a mechanism of proton-driven conformational shifts that regulate solvent access to the active site, ensuring that only the substrates of the hydration reaction can enter.

## Materials and Methods

### Screening thermophilic homologs

The list of DabA protein IDs from *Desmarais et al*. (22) was cross-referenced with the TEMPURA database (45) to find thermophilic homologs of the complex. Five candidates were chosen and the Dab operons were codon optimized and ordered as gBlocks from Twist Biosciences (Supplementary Table 5). The homologs were cloned into a pFE backbone (see below) with the Dab operon downstream of the Tet promoter (46), and the resulting plasmids were transformed into CAFree and plated on LB plates containing 100 nM anhydrotetracycline (aTc). Homologs that rescued CAFree growth in atmospheric air were tagged with a 9xHis and Twin-Strep tags, and their expression levels were assessed using western blotting. The *T. albus* Dab1 operon was chosen on the basis of its robust rescue of CAFree and relative lack of degradation when heterologously expressed.

### Cloning DAB constructs

All cloning was performed using either Q5 (NEB #M0491) or PrimeSTAR^®^ Max (Takara R045A) polymerase for DNA amplification and Golden Gate Cloning (47) with BsaI-HF^®^v2 for assembly. Primers were purchased from Integrated DNA technologies and are listed in Supplementary Table 3. All plasmids were sequence confirmed using nanopore plasmid sequencing. Important plasmids for this study are listed in Supplementary Table 4.

### Analyzing Sequence Conservation

To analyze conservation, we started with the ∼200 sequences in the DabA1 cluster in the UniRef50 Database. We removed any identical sequences and manually filtered for sequences that were confirmed to contain DabC in their operon, to remove any DabA2 sequences. 50 diverse DabA1 sequences were chosen for MSA and conservation analysis (Supplementary table 6). Results were analyzed using Jalview (48) and figure panels showing sequence conservation were generated using WebLogo 3 (49).

### Protein Expression and Purification

BL21-AI *E. coli* were transformed with pNRP048 and plated on LB agar plates containing kanamycin and grown at 37°C overnight. Individual colonies were picked into 5 mL of LB broth and grown, shaking, at 37°C overnight, then diluted into 1L of TB broth (RPI, SKU: T15100) supplemented with 4 mL of glycerol and kanamycin. Cultures were grown at 37°C, shaking at 180 rpm, until the OD reached ∼0.6, at which point Dab expression was induced with aTc and ZnCl_2_ added to a final concentration of 100 nM and 10 μM respectively. The cultures were grown for an additional 3 hours at 37°C, then the cell pellets were collected by centrifugation at 3500 rcf for 30 minutes, flash frozen in liquid nitrogen, and stored at −80°C until lysis.

Cell pellets were thawed on ice and resuspended using lysis buffer (50 mM HEPES pH = 7.5, 300 mM NaCl, 10 μΜ ZnCl_2_, 1 mM PMSF, 0.1 mg/mL lysozyme supplemented with 1 tablet of cOmplete^™^ EDTA-free Protease Inhibitor Cocktail (Roche, COEDTAF-RO) per 2 L of cells lysed) at a ratio of roughly 5 mL of buffer per 1 g of cell pellet. The cells were lysed in an ice bath using a Qsonica Q500 sonicator alternating 5 seconds on at an amplitude of 70% with 10 seconds off for a total on time of 7.5 minutes. Lysate was clarified at 12,000 rcf for 30 minutes, then the membrane fraction was collected from the supernatant by ultracentrifugation at 150,000 rcf for 90 minutes.

The supernatant was discarded and the membrane pellets were gently rinsed using a serological pipette with ice cold HBSZ buffer (50 mM HEPES, 300 mM NaCl, 10 μM ZnCl_2_, pH = 7.5), and then resuspended using a paintbrush in solubilization buffer (50 mM HEPES, 300 mM NaCl, 10 μM ZnCl_2_, 1mM PMSF, 5 mM TCEP, 1% w/v LMNG (anatrace, NG, 310), pH = 7.5) at a ratio of ∼5 mL solubilization buffer per gram of membrane pellet. After resuspension, small stir bars were used to mix the suspension at ∼240 rpm at 4°C for one hour.

The suspension of solubilized membrane pellets was clarified at 33,000 rcf for 1 hour, and the supernatant was mixed with Ni-NTA resin equilibrated in HBSZ buffer and incubated at 4°C for 90 minutes with gentle rocking. The mixture of resin and supernatant was then transferred to a gravity flow column and washed with Ni wash buffer (50 mM Hepes, 300 mM NaCl, 20 mM imidazole, 10 μM ZnCl_2_, 0.05% w/v LMNG, pH = 7.5) until the A280 was equal to that of the background. The complex was then eluted using Ni elution buffer (50 mM Hepes, 300 mM NaCl, 250 mM imidazole, 10 μM ZnCl_2_, 0.05% w/v LMNG, pH = 7.5) in fractions of 1 CV until the A280 was equal to background.

Each eluted fraction was heat treated at 70°C for one hour to precipitate contaminating *E. coli* proteins, then clarified at 21,300 rcf for 10 minutes, and analyzed using SDS-PAGE. Fractions containing the DAB1 complex were pooled and concentrated using 100 kDa MWCO centrifugal filters (Fisher, UFC9100) at 3,500 rcf to a volume of ∼300 μL. This concentrated protein solution was clarified again at 21,300 rcf for 10 minutes then loaded onto a Superose 6 Increase 10/300 GL (Cytiva, 29-0915-96) equilibrated with SEC buffer (50 mM Hepes, 300 mM NaCl, 10 μM ZnCl_2_, 0.001% w/v LMNG, pH = 7.5). Each fraction of interest was analyzed using SDS-PAGE. Small samples of the most concentrated fraction of each peak found to contain the DAB1 complex were flash frozen in liquid nitrogen, and then every fraction in a peak was pooled, concentrated at 10,000 rcf using Pierce™ 0.5 mL 100 kDa MWCO Protein Concentrators (ThermoFisher, 88503), flash frozen, and stored at −80°C until downstream use.

### Cryo-EM Sample preparation and Data Acquisition

The samples imaged on C-Flat 2/2 400 mesh copper grids (EMS CF-224C). All grids were glow discharged in a PELCO easiGlow at 0.39 mBar and 15 mA for 35 seconds, then graphene oxide was deposited on all grids using the drop cast method (50) modified to include cleaning with chloroform and coating the grid in PEI as described in (51). 3 μL of sample at various concentrations (0.06-0.2 mg/mL) were placed on the grid, blotted, and plunge-frozen in liquid ethane using an FEI Mark IV Vitrobot set to 6°C, 100% humidity. Blot times were 3, 3.5, or 4 seconds. Grids were clipped for autoloading and stored in liquid nitrogen. Grids were screened and the most promising grid was selected for further data acquisition.

Data for single particle analysis were collected on a Krios G3i 300kV CryoTEM (ThermoFisher) equipped with the GATAN K3 Direct Electron Detector recording in super resolution CDS mode at 81,000x for a pixel size of 0.525Å. A total of 6,032 movies were acquired using Serial EM (version 4.2.19) multishot acquisition with beam/image shift in an 8×8 array, collecting four movies per hole with a defocus range set to −0.5 and −1.8. All movies consisted of 50 frames with a total dose of 50 e-/Å2.

### Cryo-EM Image Processing

All analysis was carried out using CryoSparc v4.7.0 (52). Raw movies were motion corrected using Patch Motion Correction with default parameters. Patch CTF estimation with default parameters was used to estimate the CTF of each micrograph, then only micrographs with a CTF fit resolution between 1.77 and 3.5 were retained, for a final count of 5,812 exposures chosen for further analysis.

Blob picker was used to pick an initial set of particles 85-150Å in diameter from 300 micrographs. The particles were extracted in a 576 px box, fourier cropped to 128 px (for 2.4 Å/px), extracted, and 2D-classified using default parameters. Particles contributing to classes resembling the expected structure were chosen for Ab-initio reconstruction of three different classes of particles, and the particles that contributed to the most reasonable structure were once again 2D classified, producing a more varied array of reasonable 2D classes. These particles were then supplied to template picker job on the same 300 micrographs to pick a cleaner set of particles than the initial blob picker. 126,118 particles were extracted using a box of 576 px fourier cropped to 72 px (4.2 Å/px), then classified into three classes by Heterogeneous refinement. The 55,305 particles contributing to the class that resembled a full protein complex were used as a Topaz training set (53). The model was trained using a ResNet8 architecture with default model parameters. Preprocessing parameters were set to a downsampling factor of 8, and training parameters were defaults except the following: 700 expected particles per micrograph, a learning rate of 0.0001, 20 epochs, and 10,000 iterations per epoch. Topaz extract was used on the full dataset of curated micrographs, and 3,281,776 particles were extracted with a 576 px box size, fourier cropped to 144 px (for 2.1 Å/px).

Three rounds of heterogeneous refinement were performed with default parameters with a refinement box size of 72 voxels and a spherical mask of 130 Å until a final set of 1,119,446 particles was chosen and extracted a box size of 576 pixels with no fourier cropping. This set of particles was improved using three jobs of Global CTF Refinement: the first used 2 iterations to fit anisotropic beam magnification, the second used 2 iterations to fit the beam trefoil, and the third used 2 iterations to fit the beam tilt. Non-uniform refinement (54) was performed on this set of particles and resulting volume was used as a reference volume for reference motion correction (55) of all the particles, leading to 1,060,973 particles that were fed into Non-uniform refinement to produce a volume of the Dab complex at an estimated resolution of 2.16Å (global FSC = 0.143).

Although the overall map was of high quality, the region expected to contain the active site was still poorly resolved. A mask surrounding the active site and its surrounding flexible region was constructed in ChimeraX (56), then used as a focus map in 3D classification to classify 6 classes at a filter resolution of 4Å. Particles contributing to each class were put through separate Non-uniform refinement. Most classes were still poorly refined in the focus region, but the two largest classes (built from 168,739 and 164,827 particles) showed volumes with well-resolved density in and around the active site, and the resulting volumes were used for model building. The global resolution for both classes was estimated to be 2.29Å (global FSC = 0.143).

### Model building and Refinement

After high quality maps of the two conformations were obtained, ModelAngelo (57) was used for initial model building. Both the models for state A and state B were built and refined with similar workflows using PHENIX-v-2.0-dev-6003 (58) and Coot (59) v0.9.8.96. Half maps of each map were cropped for computational convenience to a box size of of 90.30, 118.65, 133.87Å, and were density modified and sharpened using phenix.resolve_cryo_em (60) and phenix.map_sharpening (58, 61), giving final resolution estimates of 2.12 and 2.13 Å for state A and state B respectively. The models generated by ModelAngelo were fit to the improved maps using phenix.dock_in_map, then several rounds of phenix.real_space_refine (62) followed by manual building and refinement in Coot were completed. Hydrogens were added using phenix.ready_set, and waters were added using phenix.douse, followed by manual checking. Final model fits are reported in Supplementary Table 1.

### epPCR screen in CAFree

To set up the suppression screen, site directed mutagenesis was performed on conserved proton translocation residues in DabB. Mutants were transformed into CAFree and mutants found to abrogate CAFree growth in air were chosen to co-express with the DabA library. This mutant was H229A in *T. albus* DabB and H260A in *H. neapolitanus*.

Error-prone PCR was carried out on DabA using a GeneMorph II Random Mutagenesis kit (Agilent 200552). The conditions were set according to the manufacturer’s instructions to produce ∼1 mutation per kilobase for roughly 3 mutations throughout the DabA gene. As a negative control to estimate the background rate of mutation, the same PCR was carried out using the high-fidelity polymerase, Q5. Golden gate assembly was used to insert the amplicons into a pFE backbone containing the mutant DabB. After assembly, the library was cleaned up using Zymo-Spin I columns (Zymo Research Cat: C1003) and transformed into electrocompetent CAFree *E. coli*. The transformed libraries were then plated on LB agar plates containing kanamycin and 100 nM aTc, and plated in both non-selective (5% CO_2_) and selective (ambient air) conditions. The non-selective condition was used to determine the library size. Colonies from the selective condition were miniprepped and the resulting plasmid was re-transformed into CAFree to ensure phenotype was tied to a mutation in the plasmid rather than in the genome. Plasmids that rescued growth in ambient air after retransformation were sent for nanopore sequencing.

### Kinetic CA assay

A fluorescent stopped flow assay relying on the pH sensitivity of the fluorescence of pyranine at 520 nm was adapted from (40). Experiments were performed on a KinTek AutoSF-120 stopped-flow spectrophotometer equipped with a Chroma ET520/20m 25 mm Dia Mounted (Lot # 319353) bandpass filter, with the monochromator slit width set to 0.12 mm, and the drive syringes submerged in a 25°C water bath. The optimal gain for the instrument was calculated using a blank solution of 2 μM pyranine in Ni elution buffer mixed with ultra purified water. Purified enzyme in Nickel elution buffer was minimally diluted and mixed with pyranine such that concentration of pyranine in the final solution was 2 μM and would be diluted to 1 μM in the reaction cell. This solution was kept in the dark and on ice until measurements were taken. A saturated CO_2_ solution of 34 mM was prepared by constantly bubbling gaseous CO_2_ in a round bottom flask of water submerged in a 25°C water bath. For all time courses, the enzyme and CO_2_ solutions were pipetted into their respective reservoirs and drawn into the drive syringes as quickly as possible to minimize CO_2_ escape. An initial six SF shots without any measurement were performed to fully load the reaction cell, then at least four time courses were taken. Time courses were taken with 1000 measurements for the first 10 seconds and an additional 1900 measurements for the following 190 seconds. Four time courses were taken at an excitation wavelength of 415 nm to measure any pH-independent change in fluorescence over the course of the two-minute measurement, then 4-7 measurements were taken with an excitation wavelength of 460 nm to measure the pH-dependent change in pH. The syringes were washed with buffer, distilled water, and then buffer again, before loading with the same enzyme and CO_2_ solutions that lacked any pyranine to measure the background change in fluorescence over the time course. Final curves were created by averaging each set of time courses and then graphing the following function over time with window smoothing:

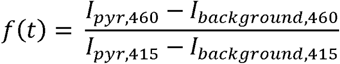

Where I is the average of the fluorescence intensity and the subscripts indicate the presence or absence of pyranine and the excitation wavelength.

## Acknowledgments

We thank Daniel Toso for cryo-EM facility support. We also thank Elizabeth Kellogg for supplying the graphene oxide and Jung-Un Park for his training on the creation of GO grids. We’d also like to acknowledge Antoine Koehl for his insightful advice throughout the project, and Professor Andreas Martin for the use of his stopped-flow instrument. This work was supported by the US Department of Energy, Physical Biosciences Program, award number DE-SC0016240 (D.F.S.). D.F.S. is an Investigator in the Howard Hughes Medical Institute.

## Notes

### Competing Interest Statement

The authors have declared no competing interest.

### Summary of Updates

We slightly expanded our discussion of possible regulation in our proposed mechanisms, and compared our structure to the recently published DAB2 structure from Halothiobacillus neapolitanus rather than comparing it to an alphafold3 model of the structure. Supplemental figures were added for this comparison.

