## Supplementary material for "Structural Insights into the Coupling Mechanism of Vectorial CO_2_ Uptake by DAB1": Supp Figs and Tables Appendix


**This PDF file includes:**

Figures S1 to S15
Tables S1 to S6
Legends for Movie S1

**Other supporting materials for this manuscript include the following:**

Movie S1

**Figures**

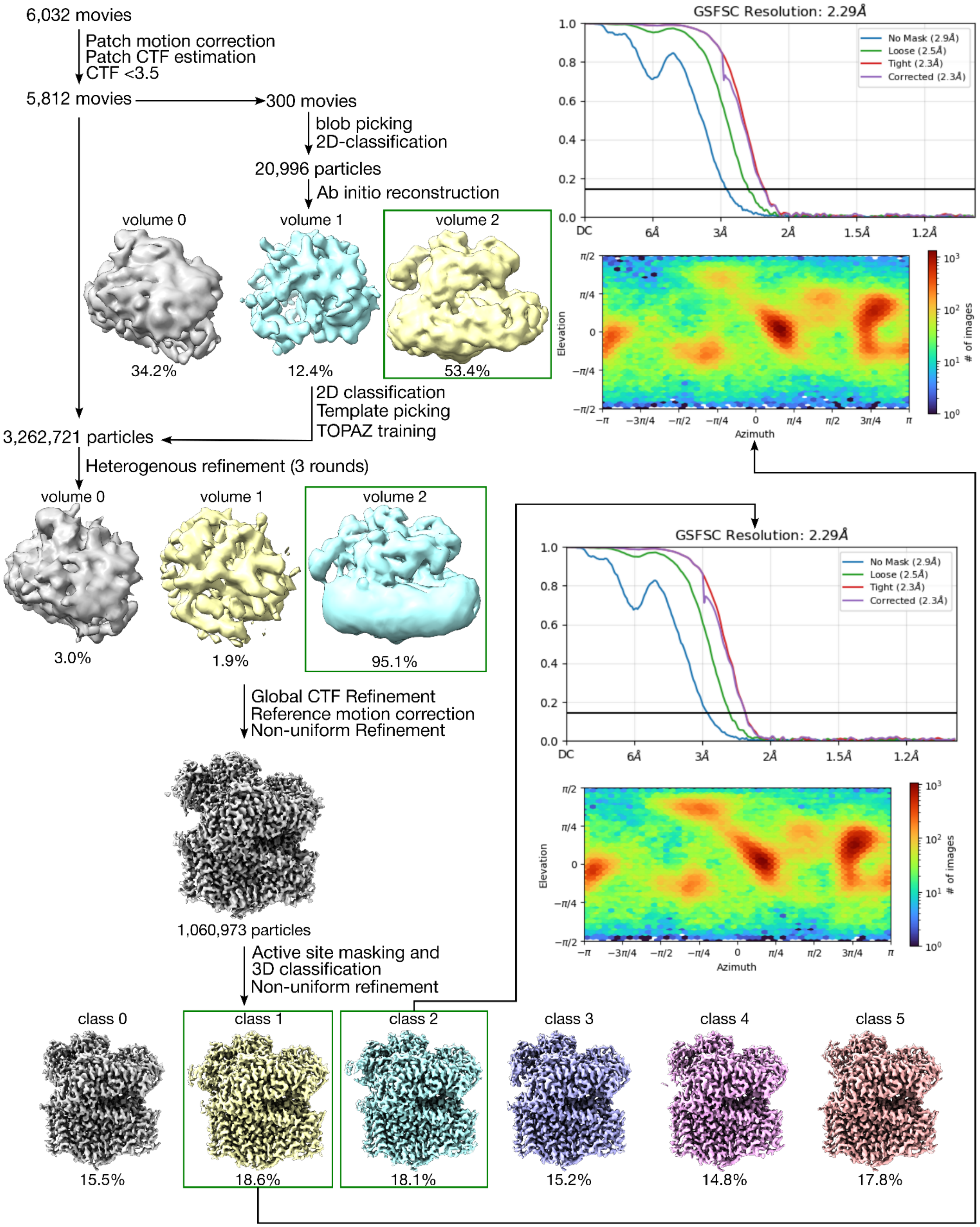


**Fig. S1. Cryo-EM Image Processing workflow.** Classes 1 and 2 were chosen for containing the best density surrounding the active site. Class 1 generated the State B model and Class 2 generated State A.

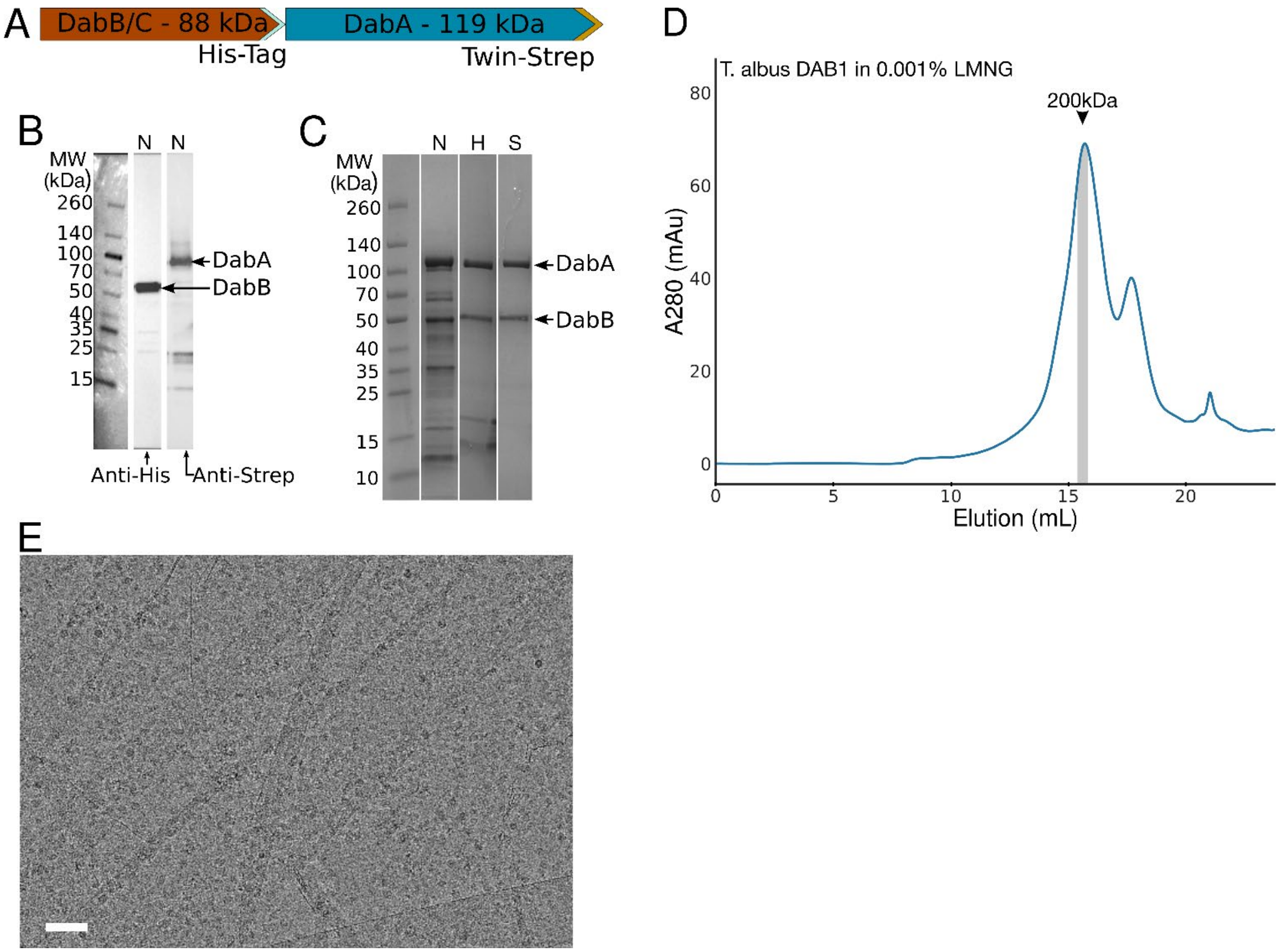


**Fig. S2. Ni-IMAC and size exclusion chromatography purification of DAB1 complex**. (A) Construct design for western blot analysis and purification of the DAB1 complex. Note that while a Twin-Strep tag was used to confirm the presence of DabA1, it was not used for affinity chromatography. (B) Western blot analysis of Ni-IMAC elution fraction confirming the presence of both DabA and DabBC after Ni-IMAC purification. N designates the fraction as a Ni elution fraction. (C) SDS-PAGE of purified enzymes after Ni-IMAC (N), 1 hour 70°C heat treatment (H) and size exclusion chromatography (S). (D) Size exclusion chromatogram of purified enzyme on a Superose 6 10/300 GL column equilibrated with 0.001% LMNG. (E) Representative micrograph of a graphene oxide grid containing purified DAB1 complex. Scale bar shows 50 nm.

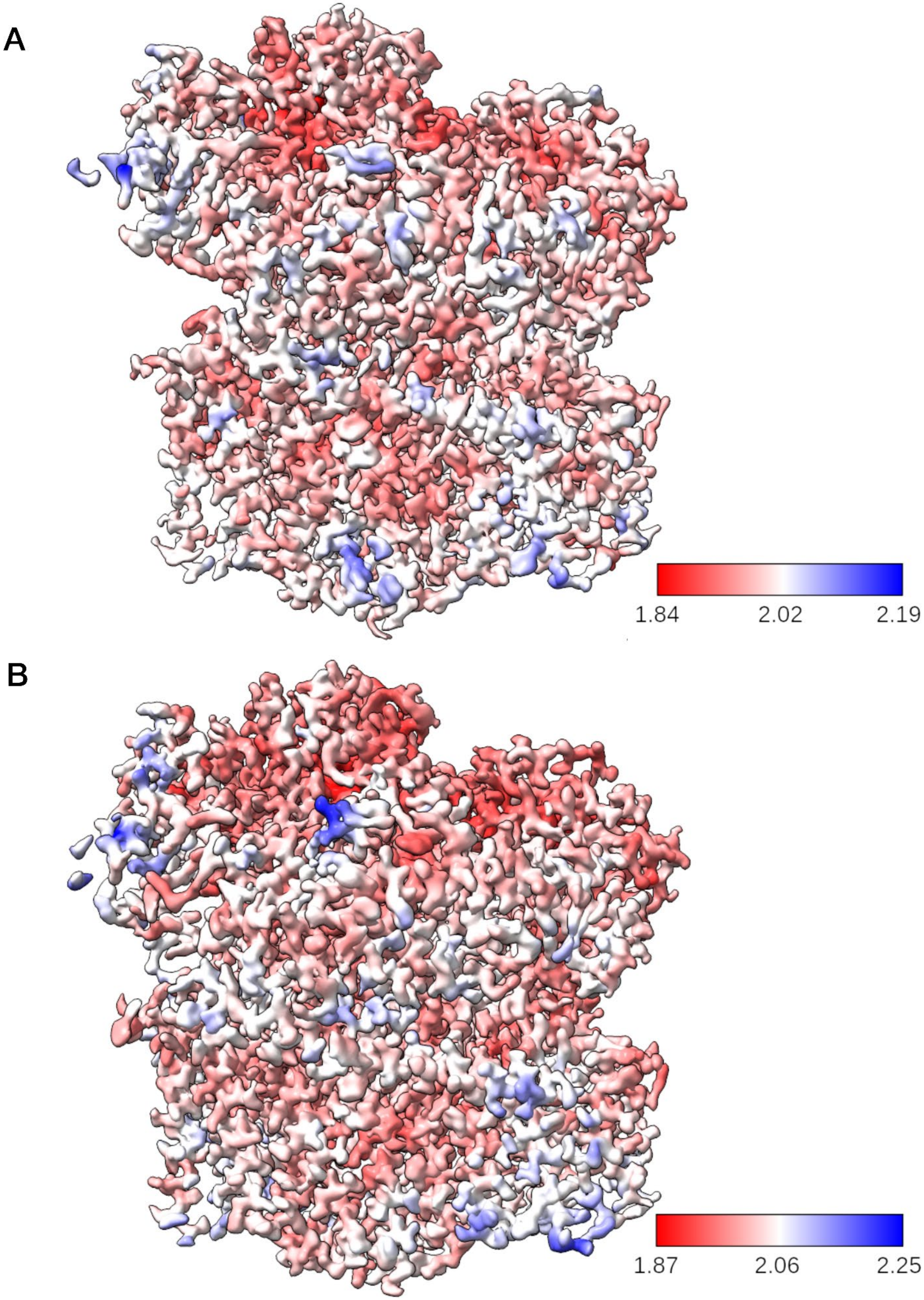


**Fig. S3. Local resolution of the two conformations of the DAB complex.** (A) State A (B) State B. Coloring generated in ChimeraX from the resolution estimates supplied by phenix.map_sharpening. Maps are shown at a threshold of 0.5.

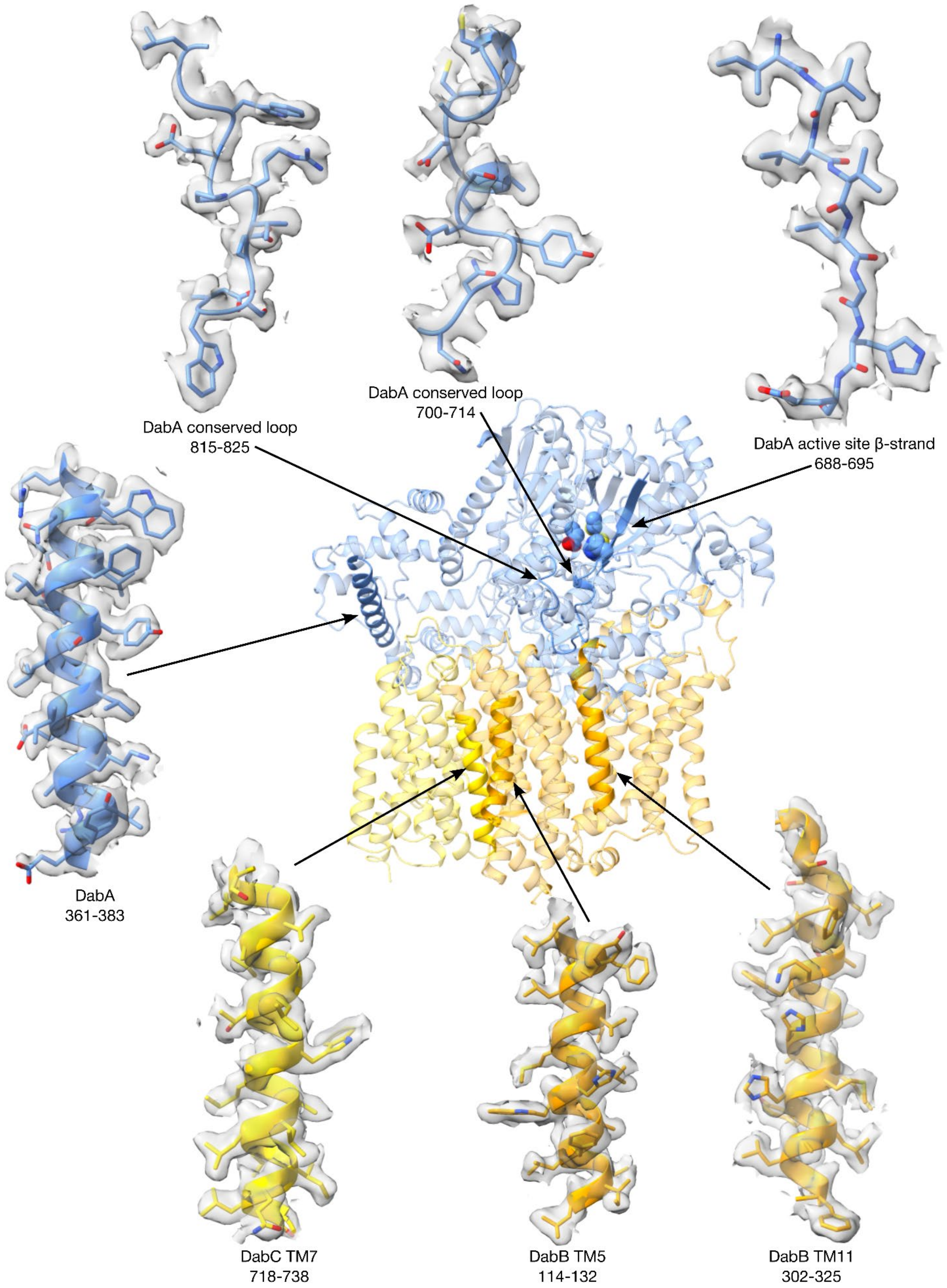


**Fig. S4. Representative density fitting in DabB and DabA.** Map and model of the State B conformation shown. Map threshold is set to 0.37.

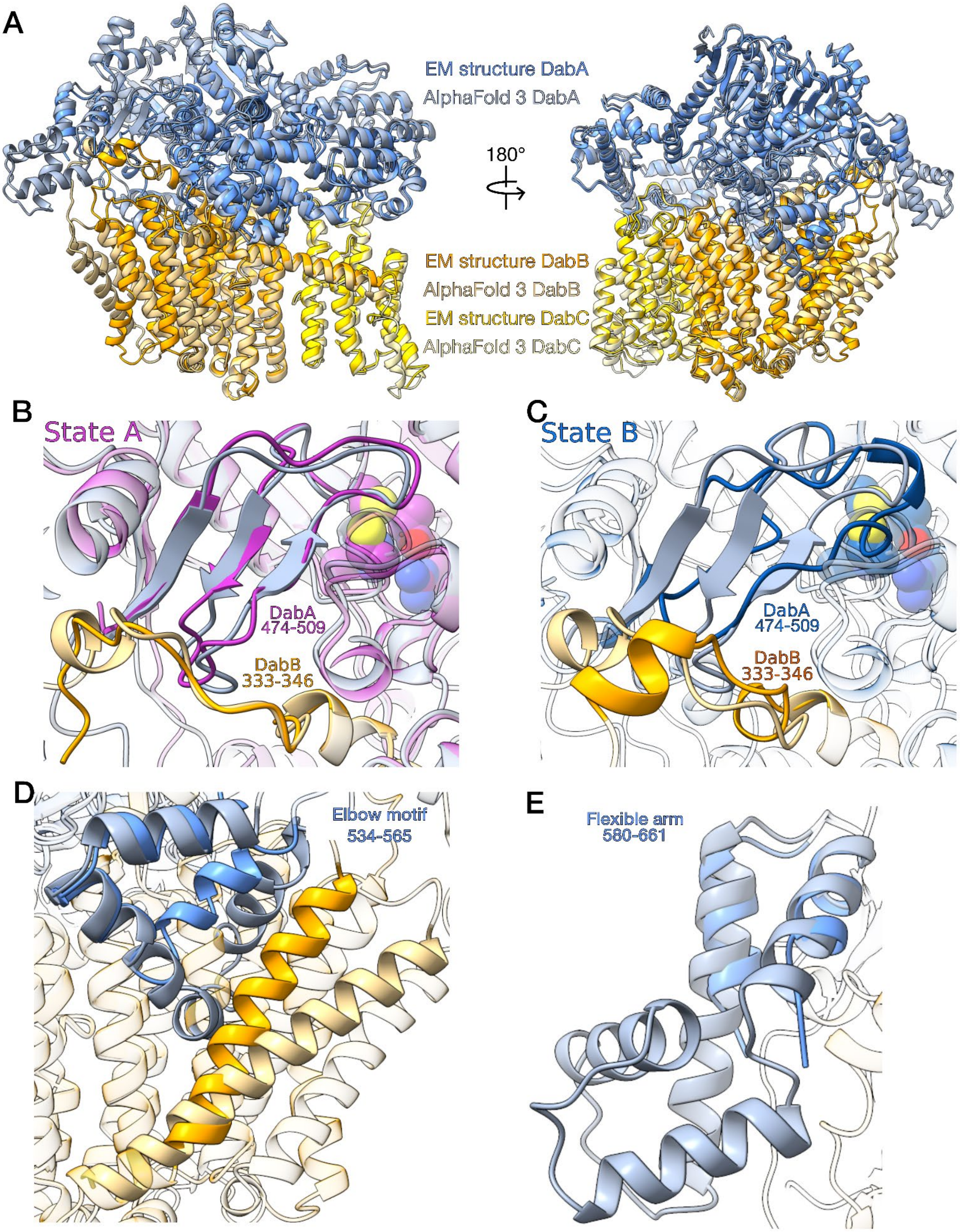


**Fig. S5. Comparison of the experimentally determined DAB1 maps with the AlphaFold3 generated model.** (A) Overlay of the two structures shown with AlphaFold3 model colored in a lighter color than the cryo-EM model. (RMSD between 524 pruned atom pairs is 0.67 Å; across all 588 pairs: 5.20 Å) (B) View of lid domain in the State A conformation and (C) State B conformation. (D) View of the elbow motif. (E) AlphaFold 3 structural prediction of the flexible arm domain that we were unable to visualize at high resolution in our maps.

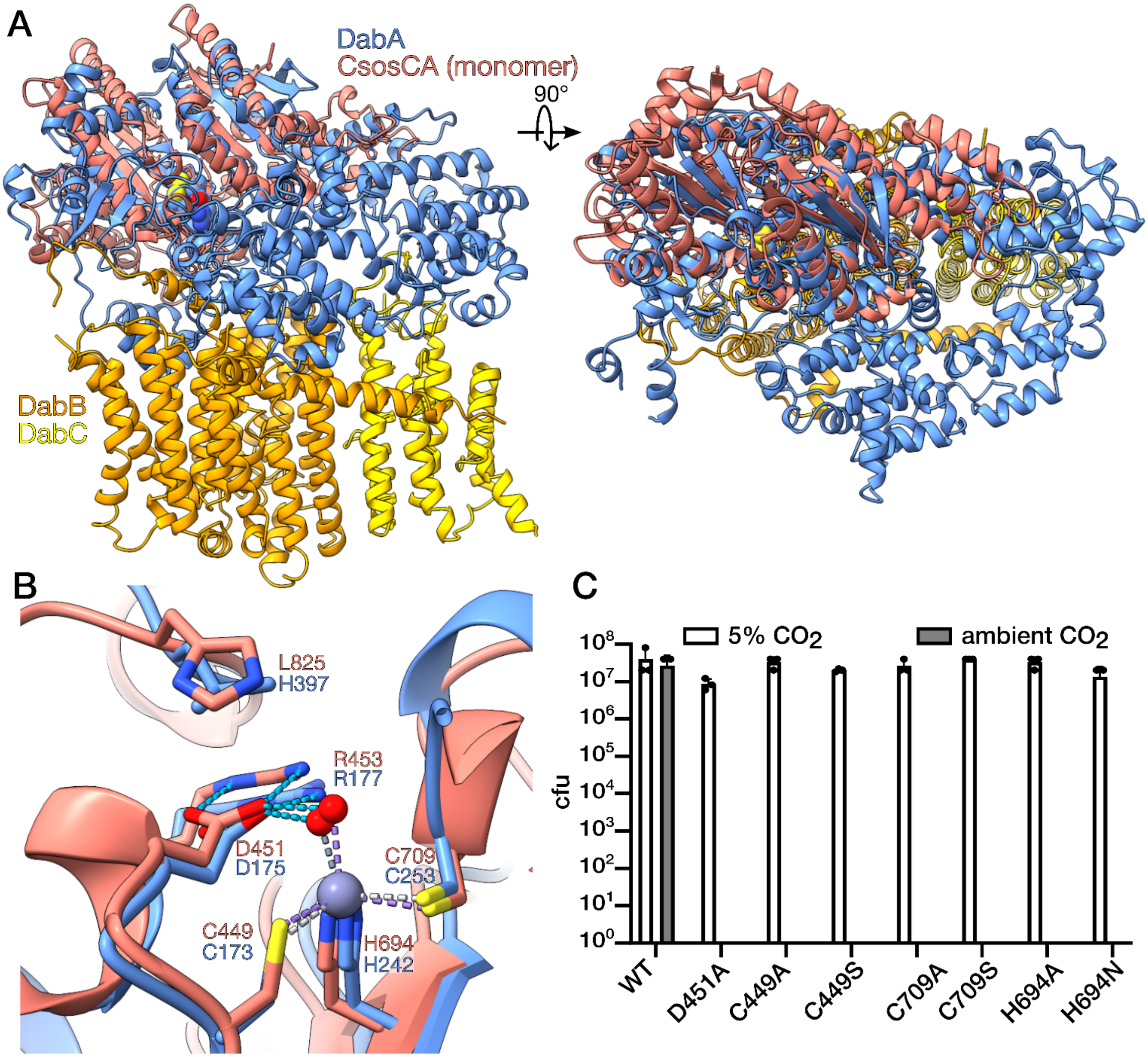


**Fig. S6. Comparison of experimental *T. albus* model with CsosCA.** (A) Structural overlay of CsosCA (PDB:2FGY) and DAB1 (State A conformation) (RMSD between 45 pruned atom pairs is 1.27 Å; across all 258 pairs: 12.55 Å) (B) View of the conserved active site structure, with the notable exception of L825. (C) Growth of CAFree expressing active site mutants in 5% $CO_2$ (permissive) and ambient air (selective). Any mutation of these residues abolishes DAB activity, and no colonies are observed in ambient air.

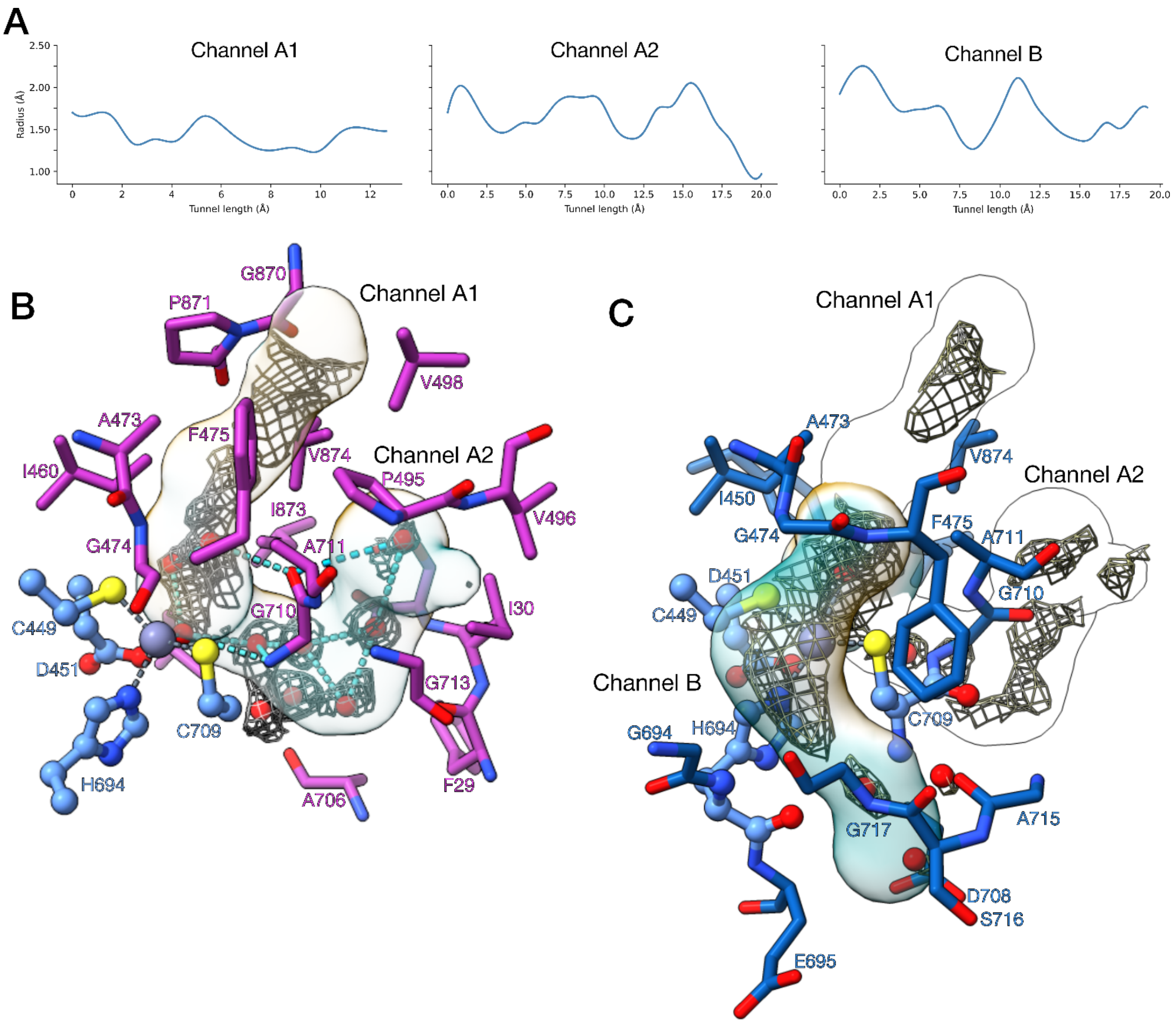


**Fig. S7. CAVER analysis of State A and State B conformations with internal electron density.** (A) Bottleneck profiles of each channel. (B) State A conformation. Both the $CO_2$ (Channel A1) and $H_2O$ (Channel A2) channels are shown with internal electron density. Channel A2 contains distinct density blobs assigned as waters while Channel A1 shows a dispersed density.  C) State B conformation. Only the hydrophilic Channel B is open, though it also contains dispersed density. Density likely from water molecules still resides in Channel A2 in this state.

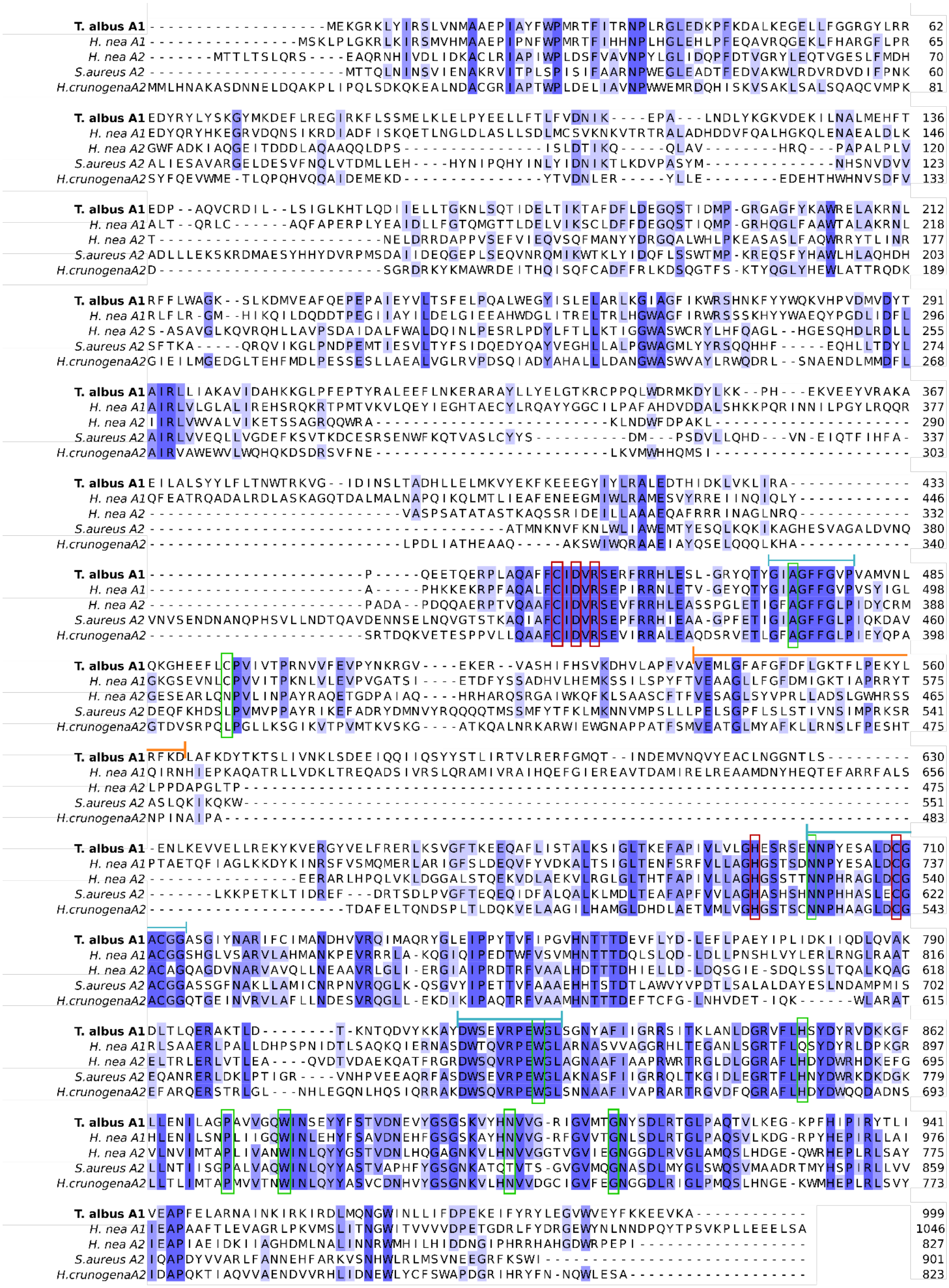


**Fig. S8. Sequence conservation of *T. albus* DabA1 compared to other DabA homologs that have been well-characterized.** Active site residues boxed in red and residues highlighted in the error-prone

PCR screen boxed in green. Elbow motif indicated by an orange bracket. Highly conserved motifs in DabA1 homologs indicated by blue brackets.

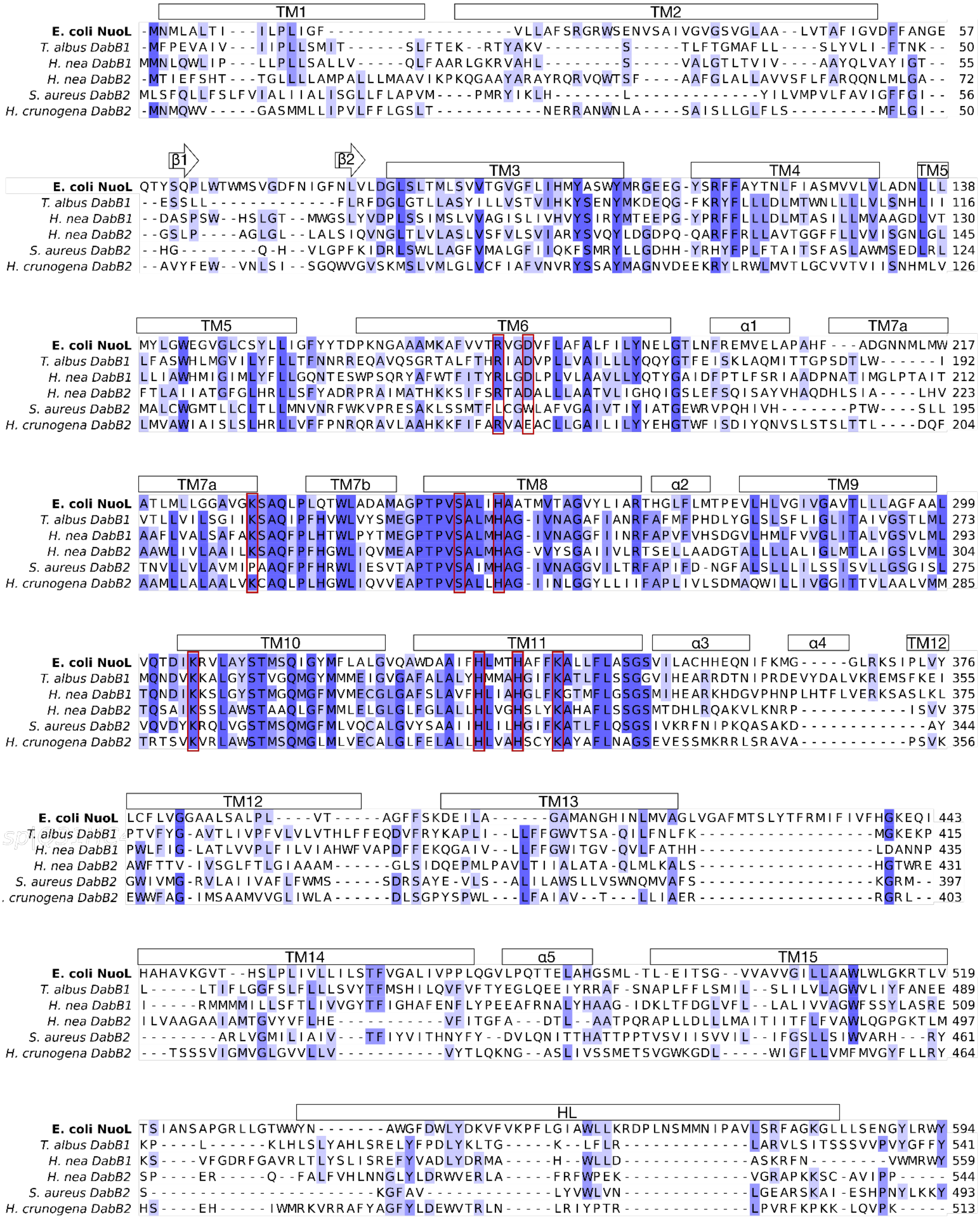


**Fig. S9. Sequence conservation of *T. albus* DabB1 compared to *E. coli* NuoL and other DabB homologs that have been well-characterized.** Proton translocation residues boxed in red. Secondary structure of the *T. albus* DabB1 are shown above. HL is the lateral helix.

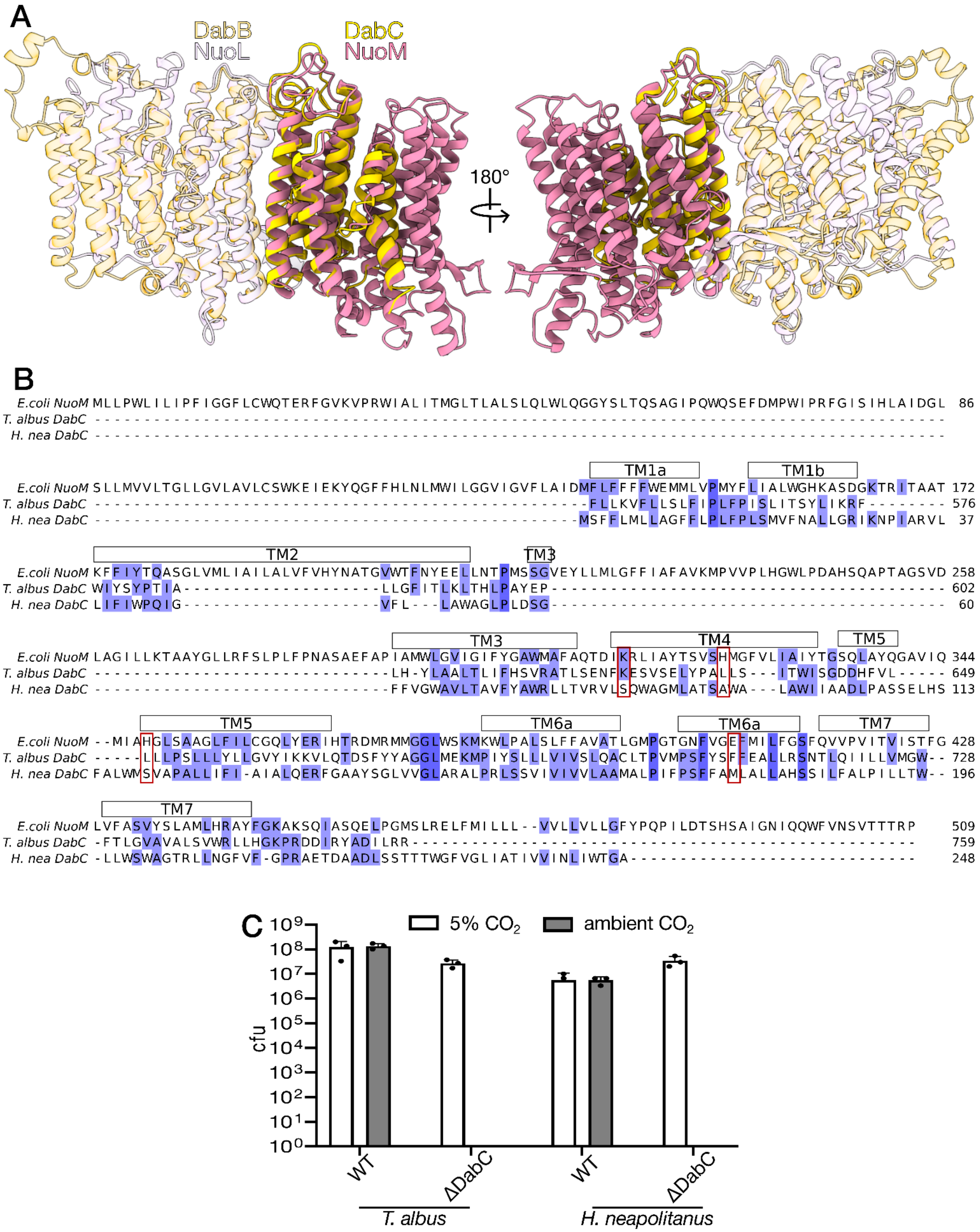


**Fig. S10. Structural and sequence homology of DabC.** (A) Structural overlay of *T. albus* DabBC and *E. coli* Complex I subunits NuoL and NuoM (PDB: 7Z7s). DabB and NuoL are shown as partially transparent while DabC and NuoM are shown in full color. (B) The MSA of *E. coli* NuoM and two DabA1 homologs shows very little sequence homology despite the structural alignment. Proton translocation residues in NuoM (boxed in red) are substituted mostly for nonpolar residues in DabA1. (C) Growth of CAFree expressing *ΔdabC* constructs of DAB1 homologs in 5% $CO_2$ (permissive) and ambient air (selective). No growth is observed in air when DabC is deleted.

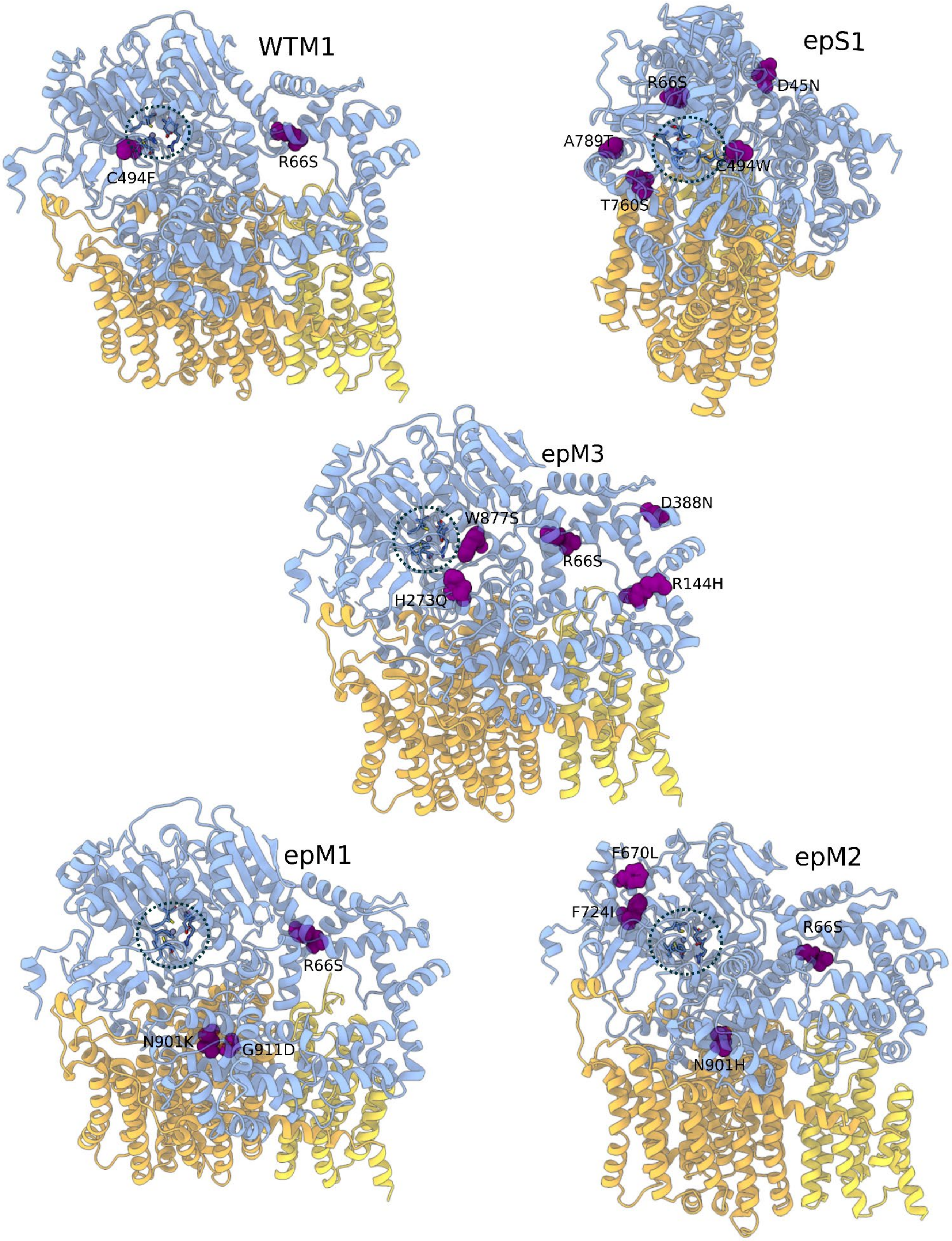


**Fig. S11. Models showing the residues mutated in variants isolated from the epPCR screen of *T. albus* Dab1.** Mutated residues shown as purple spheres in the context of the full structure. The active site is circled by the dashed line.

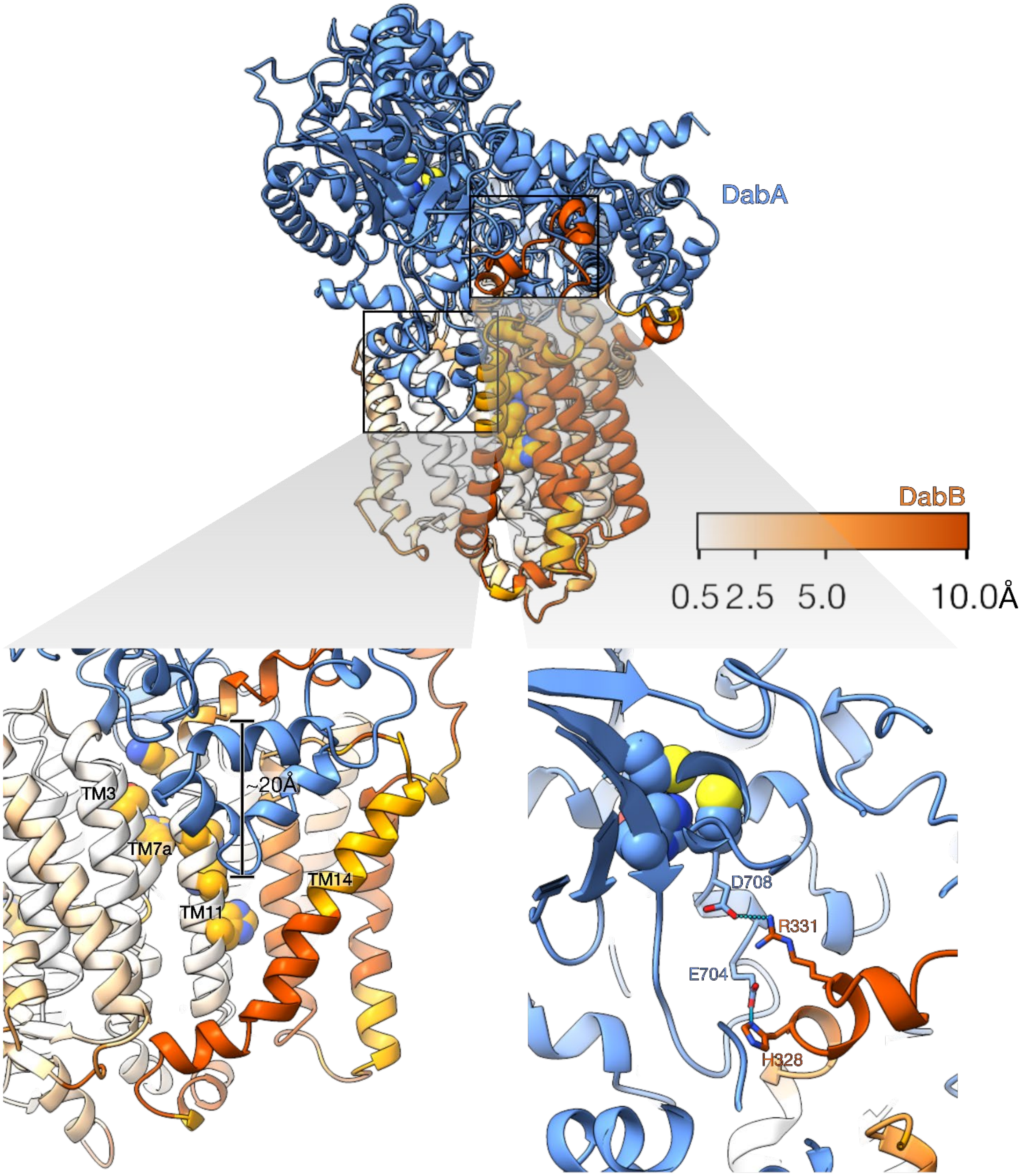


**Fig. S12. DabA and B swap helices to transmit conformational changes.** Model is State B in which DabB is colored according to RMSD value to *E. coli* NuoL. Insets show the disruption in the DabB TM helices caused by the elbow motif, and the helix DabB extends to interact with the DabA active site.

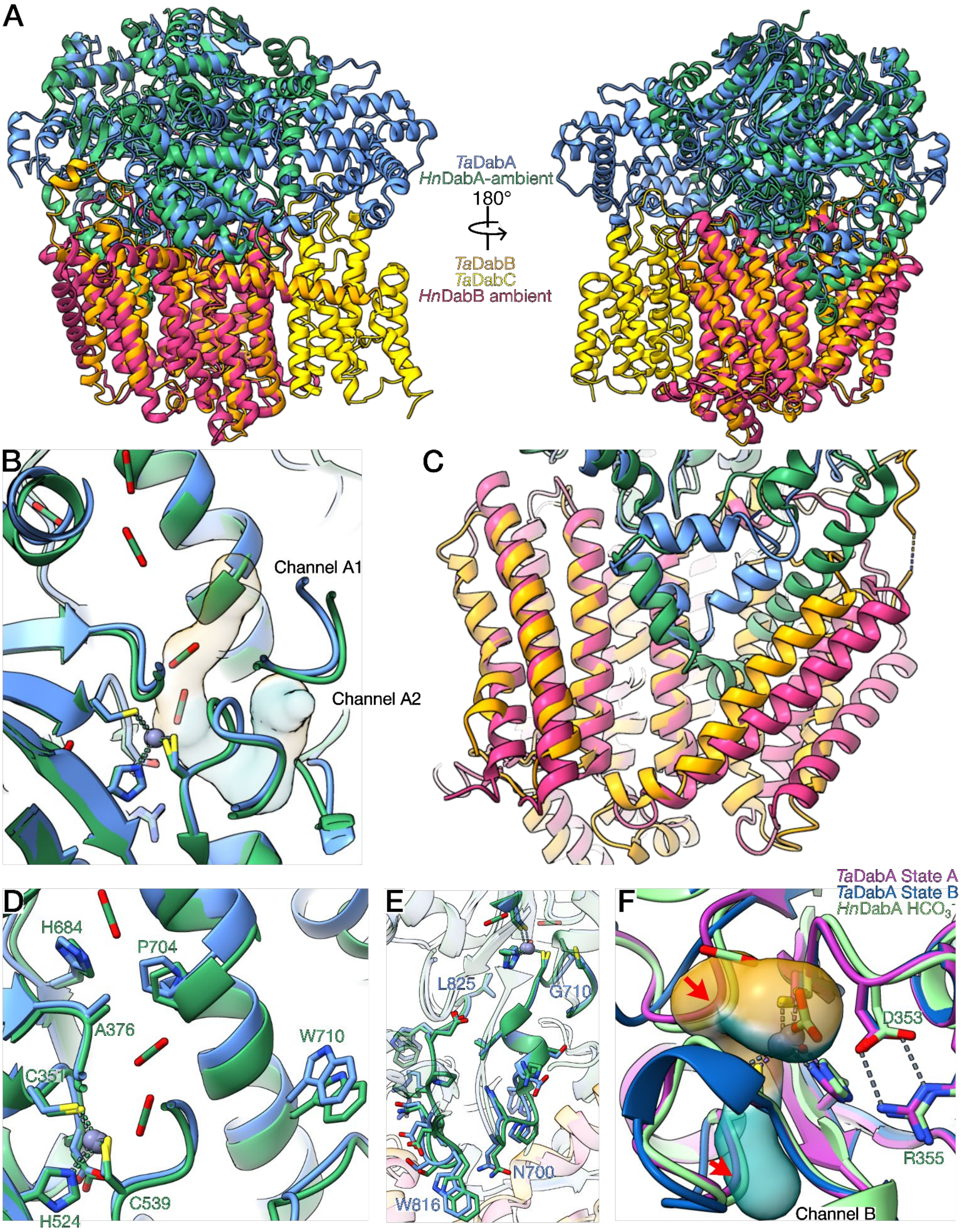


**Fig. S13. Comparison of *T. albus* DAB1 and *H. neapolitanus* DAB2 complexes.** (A) Overlay of State A of TaDAB1 complex and the HnDAB2 complex in ambient air (PDB:9RD0). The DAB1 complex is larger, containing DabC (shown in yellow) as well as additional domains in DabA which binds the cytosolic face of DabC. (B) The active site structure is similar between the two complexes, and the C1 and C2 $CO_2$ molecules observed in the HnDAB2 structure are found within the proposed $CO_2$ channel A1. (C) The elbow motif differs in shape between the two complexes: in DAB1 it does not extend as far into the membrane or the interior of DabB. (D) Key decoupling residues found in the suppressor screen align well between TaDAB1 and HnDAB2. (E) The structure of the conserved motifs between DabA and DabB is well conserved between TaDAB1 and HnDAB2. (F) Overlay of the structures of TaDabA in State A (magenta) and State B (blue) and HnDabA bound to $HCO_3^-$(PDB:9RD8, green). Even when bound to $HCO_3^-$, DAB2 more resembles State A in which the proposed $HCO_3^-$ release channel (Channel B) is closed, as indicated by red arrows.

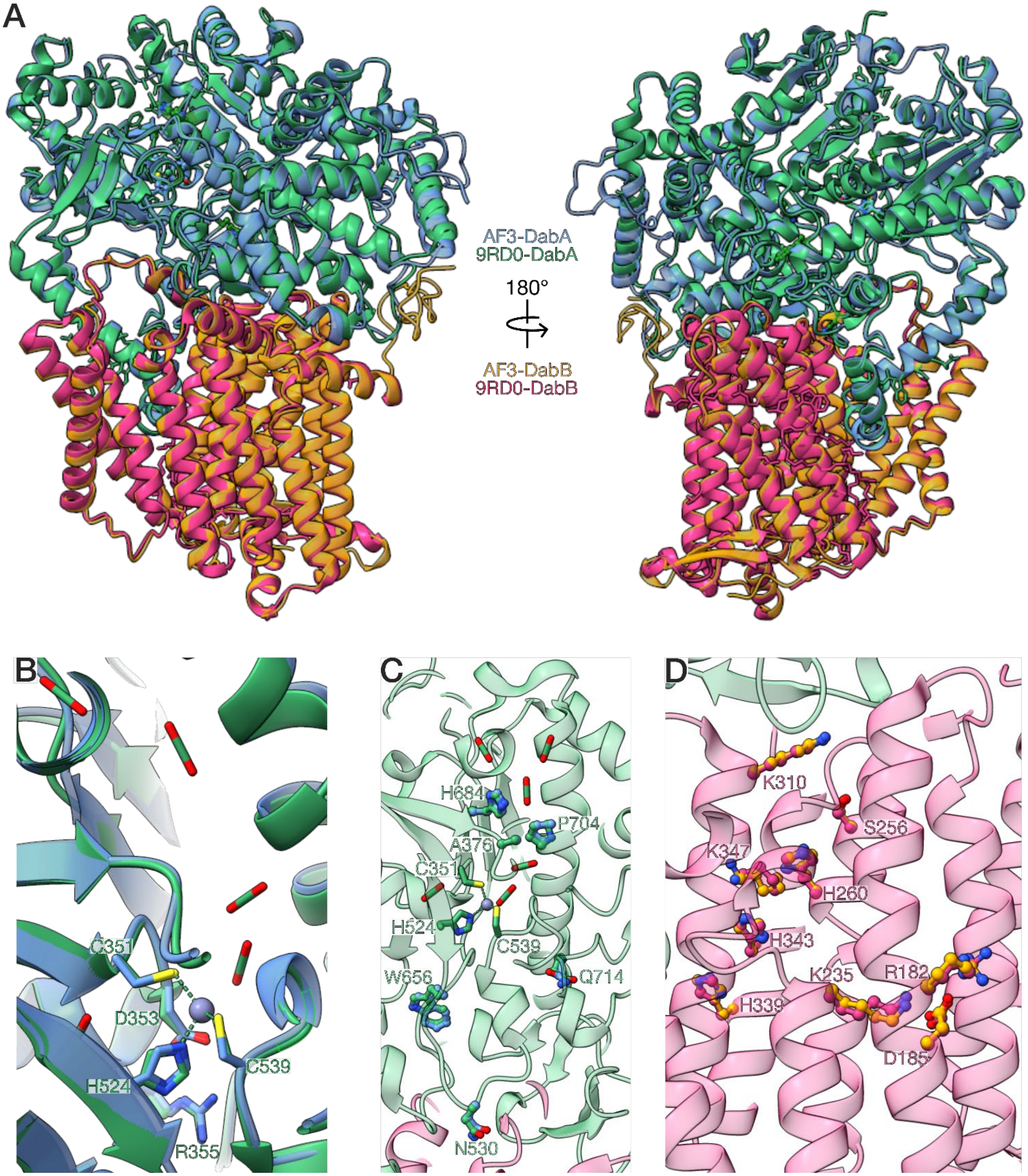


**Fig. S14. Comparison of the experimental *H. neapolitanus* DAB2 structure with the predicted AlphaFold 3 structure shows overall agreement.** (A) Overlay of the experimental DAB2 model (PDB:9RD0) and the predicted AF3 model. (B) Comparison of the active site shows excellent agreement between the predicted and experimental model. (B) The orientation of residues found to be important to coupling in our error-prone PCR screen are well-predicted by AF3. (C) The positions of conserved proton translocation residues in DabB were accurately predicted by AF3.

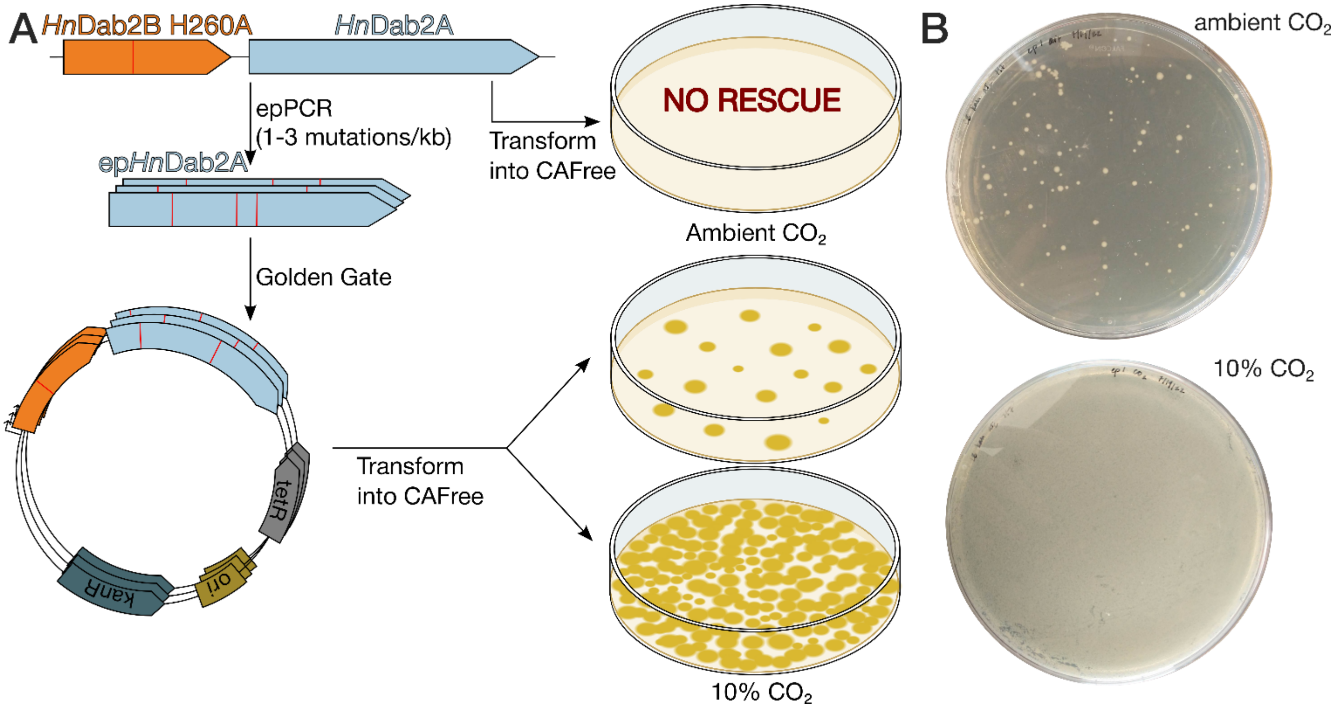


**Fig. S15. Suppressor screen performed on *H. neapolitanus* DabA.** (A) Schematic of the suppressor screen used to identify mutants in which DabA CA activity is decoupled from DabB proton translocation in the HnDAB2 complex. (B) Representative plates showing individual colonies under selective conditions (ambient air) and a lawn under permissive conditions (10% CO2).

**Tables**

**Table S1.** Cryo-EM data collection, refinement, and model building statistics.

| | **State A** | **State B** |
|---|---|---|
| **Data Collection and Processing** | | |
| Microscope | Krios G3 | Krios G3 |
| Detector | GATAN K3 | GATAN K3 |
| Nominal magnification | 81,000 | 81,000 |
| Accelerating Voltage (kV) | 300 | 300 |
| Electron exposure (e-/Å$^2$) | 50 | 50 |
| Nominal defocus range (μm) | -1.5 to -1.8 | -1.5 to -1.8 |
| Pixel size (Å) | 0.525 | 0.525 |
| Symmetry imposed | C1 | C1 |
| Initial particle images (no.) | 3,262,721 | 3,262,721 |
| Final particle images (no.) | 164,827 | 168,739 |
| Map resolution (Å) | 2.12 | 2.13 |
| FSC threshold | 0.143 | 0.143 |
| Map resolution range (Å) | 1.84-2.19 | 1.87-2.25 |
| Map sharpening B factor (Å$^2$) | phenix.map_sharpening | phenix.map_sharpening |
| **Model Composition** | | |
| Non-hydrogen atoms | 13789 | 14199 |
| Protein Residues | 1647 | 1678 |
| Water | 649 | 618 |
| $Zn^{2+}$ | 1 | 1 |
| **Model Validation** | | |
| Bonds (RMSD) | | |
| Length (Å) | 0.013 | 0.012 |
| Angles (°) | 1.51 | 1.33 |
| MolProbity score | 2.22 | 2.12 |
| Clash score | 8.15 | 7.74 |
| Ramachandran plot (%) | | |
| Outliers | 0.31 | 0.6 |
| Allowed | 4.65 | 3.54 |
| Favored | 95.05 | 95.86 |
| B-factors (mean) | | |
| Protein | 28.24 | 31.56 |
| Ligand | 44.89 | 62.98 |
| Water | 28.04 | 29.21 |
| **Model to map fit** | | |
| CC (mask) | 0.84 | 0.84 |
| CC (box) | 0.66 | 0.65 |
| CC (peaks) | 0.69 | 0.68 |
| CC (volume) | 0.81 | 0.81 |

**Table S2.** *H. neapolitanus* variants isolated with uncoupled CA activity.

| Variant | Non-synonymous Mutations present |
|---|---|
| WT1 | Q714K |
| WT2 | A376S |
| WT3 | Q714E |
| epB1 | N530D |
| epB2 | Q714K |
| epB3 | P704S |
| epS1 | P398S, L623M |
| epS2 | V652L, A659G, Q714E |
| epS3 | R258H, W656C |
| epB4 | I175F, P337A, G540C |
| epS4 | A29T, H249R, I256V, S270R, N530S |
| epS5 | Q714R |
| epS6 | H110L, W456C |
| epM1 | S178M, Q714E |
| epM2 | R622L, W656R |
| epM3 | W656R |
| epM4 | H684R |
| epM5 | G540C |

**Table S3.** Primers used in this study

| Primer number | Primer purpose | Primer name | Primer sequence |
|---|---|---|---|
| 126 | pFE backbone amplification | jjd019bbGG-F | gactgaGGTCTCtgaagcggaattcctgc |
| 127 | | jjd019bbGG-R | gactgaGGTCTCggacggtacctttctcctct |
| 128 | gblock amplification | T albus DabB F | GATCAAGGTCTCCCGTCCATG |
| 129 | | T albus DabB R | AGCTTCggtctcCATAGGGTTACCTCCGTAGAATGTCTG |
| 130 | | T albus DabA F | GATCAAGGTCTCCCTATGGAGAAAGGA |
| 131 | | T albus DabA R | GATCACggtctcGCTTCATCACGTACCGCCG |
| 128 | | A acido DabB F | GATCAAGGTCTCCCGTCCATG |
| 132 | | A acido DabB R | TCATTGggtctcCGTCACTTCAAAATCGCCTCCTTCA |
| 133 | | A acido DabA F | CAGCTAGGTCTCGTGACGC |
| 134 | | A acido DabA R | GCGCATggtctcGCTTCAACCTCTCGCGCTTG |
| 128 | | F thermo DabB F | GATCAAGGTCTCCCGTCCATG |
| 135 | | F thermo DabB R | AGCTTCggtctcCTGCATAGAATGCTCCGACGC |
| 136 | | F thermo DabA F | ATCCAGGGTCTCATGCAGACC |
| 137 | | F thermo DabA R | GATCACggtctcGCTTCATTCTAATTCCAGGACTTGCCATTG |
| 128 | | T marianensis DabB F | GATCAAGGTCTCCCGTCCATG |
| 138 | | T marianensis DabB R | AGCTTCggtctcCTTCATCGCAAAATCGCCTCCTTC |
| 139 | | T marianensis DabA F | CAGCTAGGTCTCATGAAGACCG |
| 140 | | T marianensis DabA R | GATCACggtctcGCTTCAGCCCGCACG |
| 141 | | A caldus DabB F | GATCAAGGTCTCCCGTCCGT |
| 142 | | A caldus DabB R | AGCTTCggtctcGCTCATTGTACACCTCCAGATTGACC |
| 143 | | A caldus DabA F | ACTGACGGTCTCATGAGCTCTAACAA |
| 144 | | A caldus DabA R | GATCACggtctcGCTTCAAATTGATACCCACCGTCCTT |
| 186 | cloning T. albus Dab excluding pII | Ta-nopII-pFE-gg-R | ggctacGGTCTCgcttcaTCATGCTTTCACCTCC |
| 187 | DabB proton translocation mutants | H122A_F | cgtcatGGTCTCCGCGAGCTGGgcgCTGATGGGAG |
| 188 | | H122A_R | cgtcatGGTCTCCTCGCGAACAGAATAATCAGATGGTTG |
| 189 | | D156A_F | cgtcatGGTCTCCCGCATCGCAgcgGTACCGTTAC |
| 190 | | D156A_R | cgtcatGGTCTCATGCGGTGGGTGAAAAGCG |
| 191 | | K204A_F | cgtcatGGTCTCGGGAATCATTgcgTCAGCCCAAATTC |
| 192 | | K204A_R | CGTCATggtctcATTCCCGAAAGAATAACCAGCAGAG |

| | | | |
|---|---|---|---|
| 193 | | H211A_F | CGTCATggtctcAATTCCGTTCgcgGTATGGTTAGTGTATTCC |
| 194 | | H211A_R | cgttgaGGTCTCGGAATTTGGGCTGACTTAATGATTC |
| 195 | | H229A_F | CGTCATggtctcCGCATTAATGgcgGCTGGTATTGTTAATGCCG |
| 196 | | H229A_R | CGTCATggtctcAATGCGCTTACCGGCGTC |
| 197 | | H308A_F | CGTCATggtctcGGCGCTGTATgcgATGATGGCACATGGTATC |
| 198 | | H308A_R | CGTCATggtctcAGCGCCAGAGCGAATGCGCCA |
| 199 | | H312A_F | CGTCATggtctcTATGATGGCAgcgGGTATCTTTAAAGCGACC |
| 200 | | H312A_R | CGTCATggtctcATCATATGATACAGCGCCAGAG |
| 201 | | K316A_F | CGTCATggtctcTGGTATCTTTgcgGCGACCCTGTTTC |
| 202 | | K316A_R | CGTCATggtctcATACCATGTGCCATCATATGATAC |
| 203 | | Q381A_F | CGTCATggtctcGTTCTTCGAAgcgGACGTGTTCCG |
| 204 | | Q381A_R | CGTCATggtctcAAGAACAGATGGGTTACCAGCAC |
| 205 | | D382A_F | CGTCATggtctcCTTCGAACAGgcgGTGTTCCGCTATAAG |
| 206 | | D382A_R | CGTCATggtctcTCGAAGAACAGATGGGTTACCAG |
| 283 | | R66S-F | GATACAggtctcACGCGAGGATTATtcTTATCTGTATTCAAAGGG |
| 284 | | R66S-R | GATACAggtctcTCGCGTCGCAGGTAGC |
| 207 | | C494Y-F | cagtgaGGTCTCAGGAATTCTTATACCCAGTTATCGTG |
| 208 | | C494F-F | cagtgaGGTCTCAGGAATTCTTATTCCCAGTTATCGTG |
| 209 | | C494W-F | cagtgcGGTCTCAGGAATTCTTATGGCCAGTTATCGTG |
| 210 | | C494X-R | cagtcaGGTCTCATTCCTCATGACCTTTCTGCAAG |
| 211 | error prone DabA hit single mutants | F670L-F | cagtcaGGTCTCGAGGAACAAGCTTTGCTGATCAGTAC |
| 212 | | F670L-R | cagtcaGGTCTCTTCCTCTTTGGTAAAGCCAACC |
| 213 | | F724I-F | cagtcaGGTCTCGCGCGCATTATCTGTATAATGGC |
| 214 | | F724I-R | cagtcaGGTCTCGCGCGCATTGTAAATACCG |
| 215 | | T760S-F | cagtcaGGTCTCCAATACAACAAGCGACGAGGTG |
| 216 | | T760S-R | cagtcaGGTCTCGTATTGTGCACACCGGGAATAAATAC |
| 217 | | A789T-F | cagtcaGGTCTCGATCTGCAGGTTACAAAAGATTTAACAT |
| 218 | | A789T-R | cagtcaGGTCTCCAGATCCTGTATGATCTTATCTATCAAAGGG |

| | | | |
|---|---|---|---|
| 219 | | R797C-F | cagtcaGGTCTCCATTACAAGAGTGTGCCAAGAC |
| 220 | | R797C-R | cagtcaGGTCTCGTAATGTTAAATCTTTTGCAACCTGCAG |
| 221 | | V819E-F | catgcaGGTCTCGTCCGAAGAGCGACCG |
| 222 | | V819E-R | gactcaGGTCTCTCGGACCAGTCATAGGCTTTC |
| 223 | | W877S-F | cagtcaGGTCTCGTGGGCCAATCGATTAACTCAGAG |
| 224 | | W877S-R | cagtcaGGTCTCGCCCACCACGGCCGG |
| 225 | | N901K-F | cagtcaGGTCTCGAAAGTTTATCACAAGGTTGTCGGG |
| 226 | | N901H-F | cagtcaGGTCTCGAAAGTTTATCACCATGTTGTCGGG |
| 227 | | N901X-R | cagtcaGGTCTCACTTTCGACCCAGAGCCATAAA |
| 228 | | G911D-F | cagtcaGGTCTCGAGTCATGACTGACAACTATTCTGAT |
| 229 | | G911D-R | cagtcaGGTCTCTGACTCCGATTCTCCCGAC |
| 230 | DabA Active Site mutants | C449A-F | TCAGTCggtctcACAAGCTTTCTTTgcCATTGACGTTCGG |
| 231 | | C449S-F | TCAGTCggtctcACAAGCTTTCTTTTcaATTGACGTTCGG |
| 232 | | C449X-R | TCAGTCggtctcGCTTGTGCTAACGGGC |
| 233 | | D451A-F | TCAGTCggtctcTTTGCATTGcCGTTCGGTCAGA |
| 234 | | D451N-F | TCAGTCggtctcTTTGCATTaACGTTCGGTCAGA |
| 235 | | D451L-F | TCAGTCggtctcTTTGCATTctCGTTCGGTCAGAACG |
| 236 | | D451X-R | TCAGTCggtctcTGCAAAAGAAAGCTTGTG |
| 237 | | H694A-F | TCAGTCggtctcCCTTGTTCTGGGAgcTGAGTCCCGGTC |
| 238 | | H694N-F | TCAGTCggtctcCCTTGTTCTGGGAaATGAGTCCCG |
| 239 | | H694X-R | TCAGTCggtctcACAAGGACAATCGGAGCAAA |
| 240 | | C709A-F | TCAGTCggtctcCGCCTTGGATgcCGGAGCATGC |
| 241 | | C709S-F | TCAGTCggtctcCGCCTTGGATTcaGGAGCATGCG |
| 242 | | C709X-R | TCAGTCggtctcAAGGCGCTTTCGTACG |
| 243 | | C712A-F | TCAGTCggtctcTGCGGAGCAgcCGGTGGCGC |
| 244 | | C712S-F | TCAGTCggtctcTGCGGAGCATcaGGTGGCGCGA |
| 245 | | C712X-R | TCAGTCggtctcTCCGCAATCCAAGGC |
| 285 | DabC His tag | TaDabB-His-GG-F | gatacaGGTCTCaccatcaccatcaccatcacTAACCCTATGGAGAAAGGAAGAAAGCT |
| 286 | | TaDabB-His-GG-R | gatacaGGTCTCgatggtgatggtgGCCACTAGAACCAGACCTCCGTAGAATGTCTGCAT |

**Table S4.** Plasmids used in this study

| Plasmid name | Description | Antibiotic |
|---|---|---|
| pJJD019 | Inducible *H. neapolitanus* Dab2 operon in pFE backbone | kanamycin |
| pNRP034 | inducible *T. albus* Dab1 operon including pII protein in pFE backbone | kanamycin |
| pNRP035 | inducible *T. marianensis* Dab operon in pFE backbone | kanamycin |
| pNRP036 | inducible *A. acidocaldarius* Dab operon in pFE backbone | kanamycin |
| pNRP037 | inducible *F. thermophila* Dab operon in pFE backbone | kanamycin |
| pNRP038 | inducible *A. caldus* Dab operon in pFE backbone | kanamycin |
| pNRP048 | inducible *T. albus* Dab1 with a His tag on DabC and a Twin Strep tag on DabA in a pFE backbone | kanamycin |
| pNRP050 | inducible *T. albus* Dab1 with no tags and no pII protein in pFE backbone | kanamycin |

**Table S5.** TWIST gBlocks of codon-optimized DAB homologs.

| Name | Insert Length | Insert Sequence |
|---|---|---|
| *Thermo-crinis albus* DabA | 3383 | GATCAAGGTCTCCCTATGGAGAAAGGAAGAAAGCTATACATAAGGT<br>CCCTTGTAAATATGGCCGCTGAGCCAATTGCCTATTTCTGGCCCAT<br>GAGAACCTTTATCACACGAAACCCTCTGCGCGGTTTAGAGGATAAG<br>CCTTTCAAAGACGCTCTTAAGGAAGGCGAACTCCTGTTCGGTGGG<br>CGTGGCTACCTGCGACGCGAGGATTATCGTTATCTGTATTCAAAGG<br>GTTATATGAAAGACGAGTTCTTGCGCGAAGGAATTCGTAAGTTCCT<br>CAGCTCGATGGAACTGAAGCTTGAACTGCCATATGAGGAGCTCTTG<br>TTCACATTATTCGTGGACAATATCAAGGAACCCGCGTTGAACGACC<br>TTTATAAGGGCAAGGTGGATGAAAAGATCCTTAATGCACTGATGGA<br>GCATTTCACGGAAGATCCAGCACAGGTATGTCGTGACATACTGCTG<br>TCTATTGGCCTCAAACACACCTTACAAGATATAATTGAGTTACTTAC<br>GGGCAAGAATTTAAGCCAGACAATTGACGAGTTAACAATTAAAACG<br>GCCTTCGACTTTCTTGATGAAGGGCAAAGCACCATTGACATGCCGG<br>GTCGTGGCGCGGGGTTCTATAAAGCATGGCGGGAACTCGCAAAGC<br>GGAACCTGCGATTCTTTCTCTGGGCGGGGAAATCTCTCAAGGACAT<br>GGTGGAAGCATTTCAGGAACCAGAGCCAGCCATCGAATACGTACT<br>CACCAGTTTCGAGTTGCCGCAGGCACTGTGGGAAGGCTATATTAG<br>CTTGGAGCTGGCGCGTTTGAAGGGAATCGCGGGATTCATCAAATG<br>GCGTTCGCATAATAAATTTTACTATTGGCAGAAGGTACATCCAGTC<br>GATATGGTTGACTATACGGCTATCCGCTTGCTGATCGCGAAAGCAG<br>TCATTGACGCTCATAAGAAAGGGCTGCCGTTCGAGCCAACATATCG<br>TGCCCTCGAAGAATTCCTTAATAAGGAAAGAGCTCGGGCATACCTG<br>CTGTATGAGTTGGGTACTAAGCGCTGTCCACCGCAATTATGGGATC<br>GGATGAAAGACTATTTGAAGAAACCCCATGAGAAGGTCGAAGAGTA<br>CGTTCGTGCCAAAGCGGAGATCCTTGCGCTTAGTTATTATTTATTCC<br>TGACGAATTGGACCCGGAAGGTTGGCATTGATATTAATTCTTTGAC<br>TGCCGACCATCTGTTAGAGCTGATGAAGGTCTATGAGAAATTTAAG<br>GAAGAAGAAGGCTATATTTATTTACGTGCGCTGGAAGACACGCATA<br>TCGACAAACTCGTGAAACTTATCCGTGCGCCACAGGAAGAAACGCA<br>GGAGCGCCCGTTAGCACAAGCTTTCTTTTGCATTGACGTTCGGTCA<br>GAACGCTTTCGCCGGCACCTGGAGTCTCTTGGGCGATACCAGACA<br>TATGGCATCGCAGGGTTCTTCGGAGTTCCGGTGGCAATGGTTAACT<br>TGCAGAAAGGTCATGAGGAATTCTTATGCCCAGTTATCGTGACCCC<br>GCGTAACGTAGTCTTCGAGGTCCCGTATAATAAGCGTGGCGTGGA<br>GAAAGAGCGGGTCGCTAGCCACATCTTTCACTCCGTCAAGGACCA<br>TGTTTTAGCTCCGTTCGTAGCTGTTGAGATGTTAGGTTTCGCGTTC<br>GGGTTCGACTTTCTGGGTAAAACGTTTCTCCCTGAAAAGTACTTAC<br>GGTTCAAAGACTTGGCATTCAAGGACTATACCAAGACCAGCTTGAT<br>TGTTAATAAACTGTCAGACGAGGAGATTCAACAAATCATTCAGAGTT<br>ATTACAGTACTCTGATTCGTACAGTTCTGCGGGAGCGTTTCGGCAT<br>GCAGACAATTAATGACGAAATGGTAAACCAAGTCTATGAGGCGTGT<br>CTGAACGGTGGTAATACCCTGAGTGAGAATTTAAAAGAGGTCGTGG<br>AGCTGCTGCGCGAAAAGTATAAGGTTGAACGGGGATACGTTGAGT<br>TGTTTCGGGAGCGCTTGAAGTCGGTTGGCTTTACCAAAGAGGAACA<br>AGCTTTTCTGATCAGTACCGCGCTGAAGTCAATCGGGCTTACGAAG<br>GAATTTGCTCCGATTGTCCTTGTTCTGGGACATGAGTCCCGGTCTG<br>AGAATAATCCGTACGAAAGCGCCTTGGATTGCGGAGCATGCGGTG<br>GCGCGAGCGGTATTTACAATGCGCGCATTTTCTGTATAATGGCAAA<br>CGACCATGTTGTTCGTCAAATTATGGCGCAGCGATATGGTCTTGAA<br>ATCCCGCCGTACACCGTATTTATTCCCGGTGTGCACAATACAACAA<br>CCGACGAGGTGTTTCTGTACGACTTAGAATTCCTGCCGGCTGAGTA<br>TATCCCTTTGATAGATAAGATCATACAGGATCTGCAGGTTGCAAAA<br>GATTTAACATTACAAGAGCGTGCCAAGACCCTGGACACGAAGAATA |

| | | |
|---|---|---|
| | | CCCAAGACGTTTATAAGAAAGCCTATGACTGGTCCGAAGTGCGACC<br>GGAGTGGGGCCTGAGTGGGAATTATGCATTTATTATTGGTCGTCGA<br>AGTATCACTAAGTTGGCTAACTTGGATGGCCGGGTATTTCTGCATT<br>CCTATGATTATAGAGTAGACAAGAAGGGATTCCTTTTGGAGAACATT<br>CTGGCTGGCCCGGCCGTGGTGGGCCAATGGATTAACTCAGAGTAT<br>TATTTTAGCACTGTTGATAATGAGGTTTATGGCTCTGGGTCGAAAGT<br>TTATCACAATGTTGTCGGGAGAATCGGAGTCATGACTGGCAACTAT<br>TCTGATTTACGCACCGGGCTGCCGGCTCAGACTGTTCTTAAGGAAG<br>GAAAGCCGTTTCATATCCCGATCCGTTATACTTTGATTGTTGAGGCA<br>CCTTTCGAACTGGCCCGCAACGCAATAAATAAGATCCGGAAAATCA<br>GAGACTTAATGCAGAATGGATGGATCAACCTTTTAATTTTCGACCCC<br>GAAAAGGAGATTTTCTACCGCTACTTAGAAGGTGTTTGGGTTGAAT<br>ACTTCAAAAAAGAGGAGGTGAAAGCATGAAAAGTGTAAAACTCTAT<br>GAGATGAAGAAGGTGGAAATAATCGTTAGAGGCGAAGATCTGGATT<br>TCGTGCTGGACCTGCTGGATCGCGCCGGAGCAACAGGTTATACAA<br>TTATCCACAATCTTAGTGGGAAGGGTTCCCATGGATTCCATGAGGG<br>GCATCTGTTGTTCAACGAGGAAGATACACTGGTGATGGTTATATCG<br>GTGATGCCCGAAAACTTGGTTGAGGCCGTACTCGAGGGCATAACA<br>CCGTTTCTGAACAAACACTCGGGAGTAGTCTTCGTGTCCTCGGTCA<br>TGGTATCTCGGGTGGGTAAATTCGGCGACGTCCCTACGAAGGCTG<br>ACGGCGGTACGTGATGAAGCCTCTGGTCAGAT |
| *Thermo-crinis albus* DabB | 2317 | GATCAAGGTCTCCCGTCCATGTTCCCAGAAGTGGCAATCGTTATCA<br>TACCGCTGTTAAGTATGATCACAAGCTTATTCACCGAGAAACGTAC<br>GTATGCCAAAGTGAGTACGCTCTTTACAGGCATGGCATTCTTACTG<br>TCATTATATGTTCTGATCTTCACGAACAAGGAGTCAAGTCTGTTATT<br>CTTACGCTTCGATGGCTTAGGGACGCTGCTGGCGTCATATATCTTA<br>TTGGTATCCACGGTCATCCACAAGTACTCTGAGAACTATATGAAGG<br>ATGAGCAGGGATTCAAGCGCTATTTTCTGCTGCTTGACTTAATGAC<br>ATGGAACTTACTCTTGCTCGTTTTGAGCAACCATCTGATTATTCTGT<br>TCGCGAGCTGGCATCTGATGGGAGTTATCCTCTATTTCCTGTTGAC<br>GTTTAACAATCGCCGGGAACAGGCCGTGCAAAGTGGCCGCACTGC<br>GCTTTTCACCCACCGCATCGCAGACGTACCGTTACTGGTTGCAATA<br>TTACTTCTGTATCAGCAATACGGTACGTTCGAGATCTCTAAGCTTGC<br>ACAAATGATTACCACCGGCCCGAGTGACACACTTTGGATTGTTACT<br>CTGCTGGTTATTCTTTCGGGAATCATTAAGTCAGCCCAAATTCCGTT<br>CCATGTATGGTTAGTGTATTCCATGGAAGGCCCGACGCCGGTAAG<br>CGCATTAATGCATGCTGGTATTGTTAATGCCGGGGCGTTCATCGCG<br>AATCGTTTCGCGTTCATGTTCCCCCATGATCTGTATGGTTTATCCCT<br>GTCTTTTCTGATCGGACTGATCACCGCAATTGTTGGTTCAACCTTGA<br>TGCTGATGCAAAATGACGTTAAGAAAGCTCTGGGTTATAGTACGGT<br>GGGGCAAATGGGCTACATGATGATGGAAATCGGGGTTGGCGCATT<br>CGCTCTGGCGCTGTATCATATGATGGCACATGGTATCTTTAAAGCG<br>ACCCTGTTTCTTAGCAGCGGTGGGGTGATCCATGAGGCCCGGCGC<br>GATACAAATATCCCTCGTGATGAGGTTTATGATGCGCTGGTGAAGC<br>GCGAGATGTCATTCAAGGAGATTCCTACGGTATTCTATGGTGCAGT<br>TACCCTGATTGTCCCCTTCGTTCTGGTGCTGGTAACCCATCTGTTC<br>TTCGAACAGGACGTGTTCCGCTATAAGGCCCCGCTGATACTGCTGT<br>TCTTCGGGTGGGTGACTAGCGCCCAAATTCTCTTCAATCTGTTTAA<br>GATGGGGAAGGAGAAGCCGCTGTTGACTATCTTTCTGGGTGGATTT<br>AGTCTGTTCTTACTGCTTTCCGTTTATACATTCATGAGTCATATATTA<br>CAAGTTTTCGTTTTCACCTATGAGGGGCTGCAAGAGGAGATTTATC<br>GGCGTGCATTCTCGAACGCCCCGTTATTCTTCCTGAGCATGATACT<br>GAGTCTTATTTTAGTGTTAGCCGGTTGGGTCCTGATTTATTTCGCTA<br>ATGAGGAGAAGCCTCTTAAGCTTCACCTTTCTCTGTATGCTCATTTG<br>TCGCGGGAGTTATATTTCCCAGACCTGTACAAGCTGACCGGCAAAT |

| | | |
|---|---|---|
| | | TATTCCTGCGTCTCGCAAGAGTGCTTTCTATTACATCATCTTCCGTT<br>GTTCCGGTGTATGGTTTCTTCTACCAAGGTGGTTCATCGGGATCTT<br>TCCTTCTCAAGGTTTTCCTCTTGAGTTTGTTCATACCGTTGTTTCCA<br>ATTTCCTTGATCACATCTTATTTGATCAAACGCTTTTGGATTTACTCA<br>TACCCGACCATTGCGTTATTAGGGTTCATTACTCTGAAACTGACTCA<br>CTTACCAGCGTACGAACCTCTGCACTACCTGGCCGCGCTCACTCT<br>GATTTTCCACAGTGTTCGTGCTACCCTGTCGGAGAATTTTAAAGAG<br>TCTGTCAGTGAGCTCTACCCGGCTCTGCTGTCAATAACATGGATCT<br>CAGGCGACGACCACTTCGTTCTGCTGCTCCTGCCGAGCCTGCTCT<br>TATACTTATTGGGAGTCTACATAAAGAAAGTTCTGCAGACAGATTCG<br>TTCTACTATGCGGGCGGGTTAATGGAGAAAATGCCGATATACAGCC<br>TGCTGCTCGTGATAGTCAGCCTGCAGGCATGTCTCACTCCCGTGAT<br>GCCGTCCTTCTATAGCTTCTTCGAGGCATTACTGCGTTCGAATACT<br>CTCCAGATTATACTCCTTGTTATGGGTTGGTTCACCTTGGGAGTTG<br>CCGTTGCATTATCAGTTTGGCGTTTGTTACACGGAAAACCACGAGA<br>CGATATACGTTATGCAGACATTCTACGGAGGTAACCCTATGCTCTG<br>GATCGAC |
| *Therm-aero-bacter marian-ensis* DabA | 2670 | CAGCTAGGTCTCATGAAGACCGCTACGACGCCACGGCAGCGGCCA<br>GGTCGCCCGGTTCATGGGCAAGGCGGCGGTCGACCAATGGTGGA<br>ACGTGTGCGTAGAGCATGTGATGCATTAGCACCGCTTTGGCCACTC<br>GCAACTTATATTGCTAGACACCCCTGGCCTGGTTTAGAGCGTATGC<br>CTTTTGCAGAAGCTTTGGATCATCTTCAACAAGTTCAAGGGGTAGA<br>CCTGCTTCCGCCCTTAGCGCTGTGTCGGGCGGCCCTCGCAAAAGG<br>TGAAATTGACCCAGCGGTTCTGGATCGTCGCTTGGCAAGATGGCT<br>GGATGACCATGTGTCACCGCAACTTAGACCTGAAGCTGAACGTGTT<br>TGTCGTGCTTTGCTGTGGCAAGAAGAAGTTCCGCGTCCTGAAGGT<br>GGGGTAAAAGCATGGGCCGCCTGGGTTGAGGAAGCGGCTCGTGC<br>GGGACGGTTTAGACATCGATCACAACGTGTTCTTCCCCGATCTGCC<br>CAACTTCGTGTTACAGCGCGCTTAGACGCCCAAATGATTCGTTGGT<br>GTAAATTGTTTCTGGACGAAGGGCAAGCCCGATGGCCATTACCATA<br>TCGTGAAGAGGGATTGTATCGCGCAGTTAGAAGACTTGTGCCATAT<br>GACCCTGCTTTGACGCGTGCAGAACGCCGACGTACAGCCGACTGG<br>CCGCTTGATGCCGAAGTGGCATTGGTTTGGGCTTTAGAGCGTCTG<br>GGTGTCGCGGACGCCGAGGTCGACGCTTATTTAGAAGCACATTTG<br>TTGGCTCTTCCAGGGTGGGCAGGTATGCTGTGGTGGCGGGGACG<br>CCAGGCTGGGGATGAAATGGCACCCCTTGTTGATTATCTTGCAGTC<br>CGTCTGGCCTTAGAATGGACTCTGTGTGCACCGGATCTGCCAGTG<br>ACAGGCCCAGCGGATCATCATGGGTCCGCAGTATTGCCATTGTTGT<br>GGGCATGGGAACGATGGGGTGGGATGACTCCGGCACGTTGGCGT<br>CGTTTGGATCCCGAGGAGCAAGAACGCCGTCTTGCGTTAGTCGAG<br>CGATTTCTGCGTATTGATCGTCGTCTTCTTTGGTTAGAAGCGTGGG<br>AAGAAACCTATGCAGCTCGACTGCGCCGCGCATTGACAGCTGGGA<br>GCCCGTCGGATGGAGATGCAAAACCCGCCCCCGCGGTCCAGTTGT<br>TGTTTTGTATAGACGTCCGTTCTGAGCCCCTTCGTCGGCATCTGGA<br>GCGTGCAGGCCCGTTTGAAACTTATGGATGCGCTGGTTTCTTTAAT<br>CTTCCAATTCGGAAACGTGAATTGGACTCGCCATACGCCCACCCGT<br>CTTGCCCCGCAATTGTAGAACCTCAGCATGAAATTGCAGAGCATGC<br>GGCCCCAGATGAATTAGCGCCTTTTAGACGTCGTCGGAATGTTCTG<br>CGGTTTGTGGGTCAATTATTTAAAACGTTAAAACAACATTTGTTAGC<br>TTCGCTGGTCCTGCCAGAGCTCTCTGGTCCCTGGTTGGGCCTGTA<br>CATGTTGGTTCAGTCTGCGTCTCCAGGGTGGGCAGGTCGTCTGTTT<br>CATAGAGCAGAGGCGGTAGTCAAACGTAAGCCACCAACTAGACTTA<br>CTCTTGAAAGAGTCGAAGCTGGTGGCAGTGCGGGGATACCAGTTG<br>GTATGCGTACCGAGGAAATGGTTCAAGCTGTTAAAAGTTTGTTCCT<br>TTCCATTGGACTTGTATCATTTGCACCACTTGTTGTTGTTTGTGGGC<br>ACCGTAGTTTATCTACGAATAATCCATACGCGGCAGCTTTAGAGTG |

| | | |
|---|---|---|
| | | TGGTGCTTGTGGTGGTGCAGCCGGTGGTTTTAATGCCCGTGTGTTT<br>GCGGCTCTTTGTAACCGGCGTGATGTTAGAGAAGGTTTGGCCCGC<br>GAGGGTTTGCGTATACCAGATGAAACCGTTTTCGTGGCTGCAGAAC<br>ATGTAACTACCTTAGACGTTCTGCAATGGGTAGATGTCCCGCCTCT<br>TACGCCTGCAGCGCAAGAAGCTTTCGCGCGTTTATTACCGGTTTTG<br>GATCAAGTTTCTCGGCGGACACGCGCAGAACGTGTTGTAAAGCTTC<br>CTCATGTTGGGGCCGTACGGGACCCACATGCGGAAGCGTGTCGTA<br>GAGCGACCGATTGGAGTGAAGTCAGACCAGAGTGGGGCCTCGCG<br>GGAAACGCGGCGTTCGTAGTAGGAAGACGAGCCTTAACGCGTCAT<br>GTGCACCTTGATGGCCGAGTATTTCTTCATTCGTATGATTGGCGAT<br>CGGATCCATATGGTGAACGTTTAGCCGCTATAGTTGCAGGTCCTGT<br>TACTGTTGGACAGTGGATAAATTTACAATATTATGCATCGACTGTAG<br>CTCCCCACGTTTACGGATCAGGTAGTAAAGCCACACAAACTGTTAC<br>AGCAGGTATTGGGGTCATGCAAGGAAATGGATCGGATCTCATGACT<br>GGTCTTCCATGGCAATCAGTTGCAGCATCTGATCGCGAGGTAGTG<br>CATGCGCCTTTGCGTCTTCTTGTAGTGATAGAAGCCCCGCGTGAGT<br>GGATACAAAGACTTTTGCAACGTGATGCCCAATTTAGACAAAAGGT<br>TCGCCATGGTTGGATAAGACTTGCTTCTGTAGATCCAGTACGGGGC<br>GAATGGATGGACTGGGAACCCTCAGCAGTTGATGTCGCACTGTCA<br>CGTGCGGGCTGAAGCCTCTGGTCAGAT |
| *Therm-aero-bacter marian-ensis* DabB | 1541 | GATCAAGGTCTCCCGTCCATGGGGTTGATGGCATGGTGGACATTA<br>GCAGGTAGTTGGGGCTTGGCCGTTGCCTCTGCAGCCGTCTGGAAC<br>TTACGTAGAATGCCCGAAGGTTACGGGCGTTTGCACGTTGGGATTA<br>TGCTGCTGCCTATAGCGGCAGCCTTGTGGGGTCTTCGAACTGCTC<br>CCGCGGTGATAGAAGTCGGACCGTGGCGTGTGGACGCTTTGGGG<br>TGGATGATAGCACTTTATGTTGCTGCTCTGAGTGCAGCAATTCAAC<br>GTTTTACTCTCCGGTATTTGCTTGGCGATAGATCGTATAGACAAGTC<br>TTTCTGTTGTTAACGTTAACTACAGGAGCGGCGGCTTTTACTTGGG<br>TATCTGATGATGTTCGATTATTTGTTTTGAGCTGGACTCTTATGGGA<br>GCGGGTTTGGTCGCTCTCGTTGCACTTAAACGCGAATGGATGCCT<br>GCTAAAGTAGTTGCATTAACAATGGCGAAGGTTTTCGCACTGTCCG<br>CCCTCGCTTGCGCAGTTGCAATGGGGTGGCTCGCTTGGGCTACTG<br>GCTCATGGCGTCTGAGCGAAGCAATGGCACATGCTGCCGCCTTGG<br>GTCGAGGACATCAAGTGGGCATAGGTGTATTACTGGTAGTCGCAG<br>CTCTGCTTCATGCAGGCCAGTGGCCTTTTCATCGCTGGTTATTAGA<br>ATCCGCAGTATCTCCTACGCCAGTCTCAGCTGTAATGCATGCTGGT<br>ATAGTGAACGCGGGTGGCTTGCTGTTAGCACGCAGTGCTCCTTTGT<br>TAGACGCTGCTGGTTCAGGTGTAGCGGTCGCGCTGCTGACAGTCG<br>CAGGTTTGTCTGCACTGTTAGGCACAGCTATTAGCTTGGTACATGT<br>TGATTATAAAAGACAACTCGTTGCTTCAACAATGGCTCAGATGGGT<br>CTGATGTTAGTGCAGGCAGCGTTAGGGGCTTATGAAGCAGCTATTG<br>CTCATTTAGTCTTACATGGTTTGTTTAAAGCCACACTTTTCCTCAGA<br>GCTGGTTCTGCAGTCCCACGTCCAGAGGAACGCGTTTATGCCCCC<br>GGCTCTGGAAGACCCGTCCACCCAGCGGTTGGAGCCTTATTTGGT<br>GGATTAGCCGGTGGCGCTTTCTGGGCTGAAGCAAGCGGGGAACCA<br>GCTCGCTTACTGTCAGCCCTGTTCTTCGGGGCAGCGGTAGCTTTA<br>GCATGGGGTCAAGTTCCGATTTATCGCGAGGGTCGCTGGCTCGGT<br>GTAGCAGCCGCCGCACTGATGGCTTTAGCCAGCGAAGGCGTACGT<br>GTTACATTAGCAGAAGGCGTGCGTGTCGCTCTGCCCCCGGGGACA<br>AGCCCATCTCCTGTTGTTGAGGGACTGGCTGCCGTTTTATTCGCGA<br>CGGGCGCAGGTCTTCTCGCCCTTGTAGCCCGTCGGCGCACTACGG<br>GCGCTTTCGTTAGACTCTATATGACTTTGGTTAGTTTAGGCGAACCT<br>AGACCTGCTGCTATGGATCGTCACCCGCGCTATCTTGCCGCTTACG<br>TGAAGGAGGCGATTTTGCGATGAAGCTCTGGATCGAC |
| *Alicyclo-bacillus* | 2574 | CAGCTAGGTCTCGTGACGCGCTCCCGTCTCGCTCCCGCTGGAACC<br>ACGTGGACGGCGTCGTCCGCTGTTTCTGCAGCGCGTCTGGCTCTG |

| | |
|---|---|
| *acido-* | GATGTAACAGCAGTGCAATGGCCTATATCTGTTTTCATTGCGCGTC |
| *caldarius* | ATCCATGGCCTGCCCTGGAAGATCTTCCATTCGCCGCGGCCATGC |
| DabA | GTAAGCTTCAAGCCCTGGGCGGTGCGCACTTATACCCGTCTATGC |
| | GTCTTTTACGTGATGCACTTCACCGTGGAGAAATCGACCCTAACAG |
| | ATTGCGTCACAGAATGGACCAATATCTCGAGGACGATGTCCCAGCC |
| | CACTTGCGGGGCGCTTTTGCGTCAGTCCAGGAAGCACTGACTCAC |
| | GAGGCTGGGGACCCAAGTGATGCATCACCGGAAGCGATGGCTCTG |
| | GCAGCAAAAGCTGCGGCAGCACCTGACTTGCCGGATGAAGCATTG |
| | CATTTCCCTCTGCCATCGGATGGAGAACGTGCACGTGTTGATGCCT |
| | TGAGTATACGTTACTTAAAACTGTTCTTGGATAGAGGACAAGCAGC |
| | TTGGAGCATGCCTCTGCGGAGAGAAGGCTTATTCCGCGCAGCACG |
| | TGCTCTGGCAGGCCGTGACCCTAGTCTTAGTCGTGCTGAGAAACG |
| | CCGTCTCGGGGAGTTACCTGACGCTTCTGACGAAGTTTTAGCATTC |
| | GGATTGGAACGGTTCGATGTTAGCCCCAGATTTGCAGCCGATTATT |
| | TTCGAGTACACTATCTGCGCCTGCCCGGTTTTGTAGGAGCTCTCAG |
| | ATTTCTGGGCCGCGAACAAGGTCTTGAGGACCGTTGGATGCTGGA |
| | TTATCTCGCCATGCGGATTATGATTGAATGGGCACTTGCCGGAGAC |
| | GCCGGTAGAACCTTCCTGCCACATCCAGATCTGGCAGGGTTGGCG |
| | ACCTCGGCAGCAAGATTGTTGGAAGCTGCACGTGGACCGGCTTCA |
| | GTGGACCTGCTGCTGCTGGCTCACAGATATCGTGTCGCAGACCGC |
| | TATGCCGTCTGGCTTGATGCTTGGGAAGAAACCCATGAAGCACGG |
| | GTTGTACGTCGTGGTTTGGCCTTGTCAAGAGAAAATACCGTCCCTA |
| | AAGCACAATTTCTTTTCTGTATTGATGTTCGTAGTGAACCCTTACGT |
| | CGTCATCTTGAAGCAGAAGGCCCATATGAAACCTTTGGCTGTGCCG |
| | GCTTCTTTAACTTACCCGTATGGACACGTTCTTTAGATAGCTCATAT |
| | GCACATCCCTCGTGCCCGGCTATTGTACGCCCTGTAGCCGAAGTC |
| | TGTGAAGAAGCGGTTGATGAAGCCGGGCTTGCAAGATCTCGCCGG |
| | TTAATGGGAGCATGGCGCACCCTGGCCCAATCGTTTAAGAAGGTG |
| | AAACAATCCGGTGCGGCAAGCTTAGCGCTGCCTGAATTGAGTGGA |
| | CCCTGGCTTGCTTTAGATGCCGTAGCTCGTACAGTGCCCCCGTTAC |
| | GGGCGTTTGCAGCGCGAGTTGCCAGAGCAACGCTTCCACAAGCCG |
| | AAACTCGTCTGGCTTTAGACCGTCGGGAAGGCCCCGCAGGCGTTC |
| | CTCTCGGACTGAGTTCTGAAGAAATGGCAAAATTTGCAGCCGATCT |
| | TATCAGAAGTATAGGACTTACACATTTCGCCCCATTTGTAGTCGTTT |
| | GTGGCCATGAAGCCCGAGTTGAGAATAATGCCCATCGGGCAGCTC |
| | TCGATTGTGGTGCATGTGGTGGCCGCAGCGGTCGTACAAATGCCC |
| | GTGCACTGGCCGCAGTACTGAATCGGGCAGATGTAAGACGTCGTC |
| | TTGCAGCAGAGCACGGTATATTTATCCCTGGGTCAACACGCTTCTT |
| | AGCGGCTGTCCATGTAACGACGACAGATGAAATTGAATGGTTGGAT |
| | GTACCACCACTGGTAGGAGAAGCACGGGTGCAATTCGAAGCACTC |
| | TCGGCGGCAGTCCGTCGTGCGGGCGAGAAAGCGGCATCAGAACG |
| | CGCCCAAGCGTTACCTGGTGCTAAGCGCGCACGTCCAACCTTAGA |
| | AGTTCGCCGACGGGCCTCTGATTGGGCAGAAGTACGACCCGAATG |
| | GGGCCTGGCCCGAAATCGGGCATTCTGGATTGGTCGTTTGGCACT |
| | GGACAGTGGCCCGGTAGCTGGTGAAGCCTTCTTACATAGTTATGAT |
| | TGGCGTCTCGACCCTGATATACGTGGCTTAAAATCTATCGTTGCGG |
| | GACCTGTCACCGTAGCACAATGGATAAATTTACAATACTACGCCAG |
| | TACAGTAGCACCGCATGCACACGGTGCAGGTTCCAAACCTACACAA |
| | ACCGCCACTTCGGGGATTGGAGTTATGCAAGGTAATGCCAGTGAC |
| | TTACTTCCTGGTTTACCTTGGCAATCAGTTGCTAGTGATGATGCGC |
| | ATCTGTATCACAGACCGATACGTCTCCTTGTGATTATTGAAGCACCA |
| | CGCCGTATGGTAGAAAGATTGTTGCGTGAGGATGAAGGCTTCCGT |
| | CGGAAAGTCGAAAACGGCTGGCTCCGTCTGCTTATATTTGACCCAG |
| | AGCGCGGGGCATGGGCTCGTGGAGAGGCAGTAATCCGTACAAGC |
| | GCGAGAGGTTGAAGCCTCTGGTCAGAT |

| | | |
|---|---|---|
| *Alicyclo-bacillus acido-caldarius* DabB | 1550 | GATCAAGGTCTCCCGTCCATGGACTTTGCTGCCGTCCATCAGGCC<br>GTTTCACTCGTATGGATGTTAGCTTGGATTGTTGCGTCAATGACAG<br>GAGCGCTGCTGTGGCCATTGAGAGAACCGAGCAGACGAATTCGCA<br>TGCATTTAGCTTCCCTGTGTCTTGTTAGTTTAGTTAGTATGTTGGGA<br>TTGGTCACAGGGGCAATTGGGGTTCGATTAGGCCCTTTACATCCGA<br>CGGGATTGGGTTGGGCAATTAGCTCATATATTTCTGCACTTGGATT<br>AGTGGTTCAAACCTTTTCAGCCCGGCATTTAGCAGGTGATGAACGA<br>TATGGTTCTTACTTCGCCTGGATGACCTGGACGCTGTTCTTCGCGT<br>CCGCTGCTTGGATGGCAGGGGACGCATGGTTCTTTGCAGCATGCT<br>GGGTTGGCATGGACGCTGGTCTTATCAGACTCATCGCTATGCAAC<br>GCCGTAGTCGCGCAGCCCAGGCAGTTGCTCGCATGACTTGGGCTC<br>GTTTGGCTCCATCGATGGCAGGTGTTTTACTGTTATGTGGGATGGC<br>CGCATGGGCAGGACGTAGCCCTTCACTGGACACCGGAATACGTGC<br>TCTTGCGGCTCATCCGCATGCTAGTTTGGTCGCGGGGTTATTACTG<br>GGTACAGTCGCACTGGCACAAGCCGGTAATTGGCCGTTTGGTCGG<br>TGGCTTTTGGATTCTGCAGTGACTCCTACCCCAGTGTCTGCGCTGA<br>TGCATGCTGGGTTAGTTAATGCTGGTGGGCTTTTACTGGCAAAATT<br>CGCACCTGTGCTGGCCGCTGCAGGCCCTTTCCCAAGAGCTCTGTT<br>ATTGGCTTTTGCATGGCTGTCAGTTGTGACGGGTACGGGTGCTATA<br>TTAGTCCAAGCAGATTATAAACGGCAACTGGTTGCCTCAACGATGG<br>CACAAATGGGACTCATGTTAATGGAATGTGCAATGGGTGCATATGC<br>CGTTGCTATAGTCCATCTCGTTTTGCATGGTATCTTTAAAGCATCCT<br>TATTTCTTAAGTCAGGCTACGCTGTACCTGCTCCTGAAGATGCACT<br>GGTCCCACCAAGTCCTCAACCTAAAAGATCTCTCTGGCCTGCGGTC<br>GCGGGAACGCTGTTCTTACTCGCTTACGTTGCACCCCATCCTGCG<br>GACGGGATGCGTCTGCTGAGCGGACTGTTACTGGCAGCTGGGTGT<br>GCCATTGCCGTCAGATTGGCAGCTGATCTCCAAGCAGGTCGTTGG<br>CTTGGACTGGCAGCGGTAGCAGCCGCAGCTGGTGCTGCTGAGGG<br>CGTGCGGGCAGCATTAGTACGGGCAATAGCAAGTCTGGGTGCATC<br>GAATGCGCATCCGGATTGGGCTTTTGCGGTATTAGCAGCTGCTTTA<br>GTTGCATTACAAGCCTTTGCTTTTGCCTGGTGGCGTTGGCGCCCAG<br>ATACCAATGGTGCTATGCTGGCGTATGCTTGGTTGGTCCACTTGAG<br>CGACCCCCGACCAGCCTCGGTAGAGGCGCAGCCCACGTCCCTGA<br>AACAACTGGTGAAGGAGGCGATTTTGAAGTGACGCTCTGGATCGA<br>C |
| *Fonti-monas thermos-phila* DabA | 2598 | ATCCAGGGTCTCATGCAGACCTCGCCGTCCTCGCCGGCCCCCGTT<br>CACGGCGCTCATTCCTGCCGATCCGCGGCGGCTGCGGCCACACC<br>GGATTCGGGCGATGTGCGGGGTATGGATCTATATCAGCGCATCGA<br>AGCAGCCTGCGAGCGGGCGTGTCAGCAGATCGCGCCGGTCTGGC<br>CGCTGGATCGCGCAATCGCCGTCAATCCACACTGGTGCCGCATCG<br>ATCGATCCGTGCGCACGGTTGCCGCACGTATGGCTGTTCTCGGTC<br>ACATCCAGGTGTTTCCACCCCGCGACATGGTGCGGCAGGCCTGGG<br>AAGAAGGCCGCATCACTGCGGCGGATCTGGCTTACGCGCTCGATA<br>CGCTACCCGCAGCGCAAACAGCCGGTCTGACTCCGTCGGGCTGC<br>GTGCAGGCTCTGGCGGCAGCGCCGGCCCTTCCGCATTTGCCGCT<br>CTTGATCGATGTGCTCGACGACGACCCGCTGCGCCACACACGGCT<br>GTCCTGGCGGCAGGCGATCACCCATCAGGTCAGCCAGACCTGTGC<br>GGCCTACTTCGATGAGCACCAAGCCGATTGGCACCCGGACCGCCG<br>GCACGGTTTGTACGCGTTCTGGCGCGATACCCTGACGCACGACCA<br>CGGTATCGGCGTGCTGATGGGCTTGCCGCATCTCGGACGCAGCCT<br>GCATGCCCTGCCCGCCACGCGGCAGGAGGCGGAGTCTTGGGTCT<br>TGCAGCGGCTGGGTCTGCCTGAATCCGTGTGGGCGGATTACTTGG<br>AGGCAGTTCTCCTCACAGTTAATGGTTGGGCCTCATGGTGTGCGTA<br>TTTAGATTGGCAAGCACGTCAAGCAGGTCAAAGAGATATGCACCTG<br>CGCGAACTGCTGGCTATACGATTAGCGTGGGGAGCAATATTACTC<br>GAGTGTAAAGAGGACGCCTCTGCTAGACGTGCCTTCGCTGCGGTT |

| | | |
|---|---|---|
| | | CAAGCCCAGTGGAATCGGGCTGATACCTTGCTTCAGAAGGCCGAA<br>GCTTTACTTCTTGTTGATGAGGTTTGGCAACTCGCTTTTGAAGCCG<br>GCTACCAACGCCGCCTCGCTCGTCAATTACGTACCGCAGCTACAC<br>CTGCCGCACCACAACAAGTAGAAGTCCAAGCTGTGTTCTGTATAGA<br>TGTCAGATCTGAGCCGGTGCGGAGAGCTCTCGAAGCGTGCTCTCC<br>AAGCATCCAAACTATTGGTTTCGCTGGTTTCTTCGGGCTCCCGGCA<br>GCGTATACTCCTTTAGCTACTCCAGCCCGGCGGCCTCAGTTACCTG<br>GTCTGTTAGCTCCGAGTGTTGAGGTTGTAGACCGCGTTGTTGCGG<br>CTCCGGGTCAATCAGGTGCACCGACACCAGACTTAACTGTCGCCG<br>CTACACATGCCCGACAACGTCGCTTTGCCTGGGGCGGCCCATGGC<br>GCGCGGCTTCTAGATGGCCGTCAGCAGCCTATTCGTTTGTAGAGG<br>CAGTAGGTGTAGGCTACCTGGGACCTCTTATGCGTTGGCTCTGGC<br>CGTCCCCCGCGGCTTGGACACATGAAGACCACTATGGACTGCCTG<br>AACGATATCGCACAATTTGTCGACCTACGTTAATTGGCCTGGACGT<br>TGAAGAAAAGGTGGAACTTGCCGCTCGCGTATTACATGCCATGGG<br>ACTGGACCGGCGGGTAGCCCCATTAGTGCTGTTGGTTGGCCATGG<br>ATCGCAATCTGTTAATAATGCACACGCAGCAACCCTTGACTGTGGG<br>GCGTGCTGCGGTCAGACCGGGGAAGTGAATGCGCGTGCACTGGC<br>TCAGCTTTTGAACGAACCGGCAGTTAGAGCGGGGCTGCGTGATAA<br>AGGCATTGTAATACCGGAAGATACTCAATTTGTAGCAGTTCTTCACA<br>ATACTACAACCGACGAAATAGAGGGATTTGACCTGGATCGCCTTCC<br>CCCAGCTGCTCGAGCTCGTTGGACCCAACTGCGGCCTGTTTTGGC<br>ACAAGCGGGCGATCGTGTAAGACGCGAACGCGCCCCTAGACTTGG<br>ACTGGATGCCCAAGCGGACGCTAGACAGCTGCTGGCCCATATGCG<br>CCAACGTGCTTCAGACGGAGCTCAAACTCGACCAGAATGGGGATT<br>AGCGGGTAACGCGGCATTCTTAATTGCTCCAAGAAGCCGTTCCCG<br>CCACTTAATGCTTGATGGGCGGTGTTTCCTGCATGATTATGATCCT<br>GCGAATGACCCAGATGGGTCAGTCCTTGAATTATTATTGACTGCAC<br>CTATGTTAGTTACTCATTGGATTAATTGGCAATATCATGCAAGTCTG<br>TGTGACCCTCAGCGCCTTGGCTCTGGAAATAAGGTTCTTCATAATG<br>TAGTAGGCGGTCATATTGGCGTTTTCGAGGGTAACGGTGGCGATTT<br>GCGAATTGGCCTCTCTAAGCAAAGTCTTCATGATGGGACTCGCTGG<br>GTACATGAACCGATCCGTTTAACTGTCGTCGTGGACGCACCGGCG<br>GCGGCAATTGAGGGTGTAATCGCTCGGCACGCTGTCGTACGACAA<br>CTTCTGGATAATGGATGGCTGCACCTTTGGCGGTATCAAGATGCTG<br>AGTTGCTTCGTTACGCAAACGGCCAATGGCAAGTCCTGGAATTAGA<br>ATGAAGCCTCTGGTCAGAT |
| *Fonti-monas thermos-phila* DabB | 1607 | GATCAAGGTCTCCCGTCCATGCTGGCCAGTGCTTTTATGCCCGCAC<br>TGGCAATGACGCTGGCCGGGTTGCTGGTATGCCGTTCGTCCGCTG<br>GCGCCAGAGCGCTGTGGCGGTGCTTTGCATGGTTGGCAGGTGTG<br>GCATTGGCGGGCGCTGTCGTGCTCCTCGGATGGCAATGGGGTCTG<br>CCGACGCAGGCTATCGATCGCGGCGCAGCATCGGCTTGGTTTGCA<br>ACCAGCCCCCTTACCGCTTCGCTGGCAGTGCTCGTGCAGTTTCTC<br>GGCGTAGTCATCGGGGCGTTTTCGTCCCGGTATCTGGAGGGCGAG<br>GCGGGCCAGCGGCGCTATGTCGCCGCGCTTTGCGGAGTGCTCGC<br>TGCCGTACAACTGCTTTTGCTCGCCAACCACTGGCTGCCGCTGATC<br>GCGGCCTGGGTCGCCGTGGGGGCGGCCTTGCAGCCGTTGCTGTG<br>TTTTTACCCGGAACGGCCGTTCGCGTTGTTGGCAGCGCACAAGAA<br>ACGTCTGGCCGACCGGCTGGCCGACCTGTTGCTGATCGGCGCCG<br>CCGGTCTTGCCTGGTGGTCTGCAGGGAGTGGCTCTTTCGATGATC<br>TTTACGCACACCTCGCCGACGCGCAGGCTTCGGCCCTGCTGTCAT<br>GGAGCGCGGTGCTGCTTGCGGGGGCGGTGATCCTGCGTACGGCC<br>CTACTGCCGGTTCACGGCTGGTTGATTCAGGTGATGGAAGCGCCA<br>ACCCCGGTATCCGCGCTATTGCATGCCGGGGTGGTCAACCTCGGT<br>GGGATCGTGTTGATCCGGCTGGCGCCGCTGGTGGAAGCGTCGAC<br>AGCGGCGCGCTGGATATTGGTCGTCTTCGGTCTGGGGACGGCGAT |

| | | |
|---|---|---|
| | | GCTCGCCGGTCTGGTGATGCTGACGCGCATCAGCATCAAGGTCAG<br>GCTTGCGTGGTCTACCGTGGCCCAGATGGGCTTCATGGTCTTGGA<br>GTGCGGGTTGGGTCTATACACGCTGGCGGCGTTGCACCTCATCGG<br>TCACTCGCTCTACAAGGCCCATACCTTCCTCGCTGCTTCGACCGTG<br>GTGCGCCAGAGTCGCCAGCGCATGATGCGCGCAGAGGCGCGAGC<br>GTCGTTGCCCAGCCTGCTGGCGGCACCGTGGGTGGCGCTGGCGA<br>TGGTTGCAGCAGCGCAAGCCCCCTTTGCGGCATCCGCTTGGCCGC<br>TGTGGTGGAGTGGCGTTCTGGCTTTGGCATGGTCGCCTTTGCTGT<br>GGCTGCCCGCGGCGCAGGCCAATGTTCCAGGTGTGTCCCTCTTTC<br>GTGTCCTCGGGGGACTGGCTCTGATCGGTGGACTGACGGCTCTGG<br>CGCAGTTGCTGCACGGGCTGCCGCTGGGGCTTGAGGATCGGTCA<br>GCGGATGCTGCCGGCATGGTTACTTTGCTCGGCATGGTAGCGATG<br>TACCTGGCATTGCTTACCTTGCAGTGGCGGCCGCGCTGGCTCGAG<br>GGCGCGCGGCGCTGGAGCTATGCCGGGTTTTATGTGGATGAGTTT<br>TATACGCGCCTGGCGTTGCGATTTTGGCCCGCCGCCTGGACACCG<br>GCGGCAGCATCAGCGGCTGCAGCACCGTCCGGTTCGCGTCAGTAA<br>CCGTTGCGTCGGAGCATTCTATGCAGCTCTGGATCGAC |
| *Acidithio-bacillus caldus* DabB | 1679 | GATCAAGGTCTCCCGTCCGTTATGCCTTCTGTCCTCGTGCTTAGCG<br>CTCTGCCCTCTGTACGTCCTACGCGTCAATCTGCGCTGGGCACTC<br>GGTACCGCGTCCGCTGGGCGGCTATCGTAGCATTGTTAGGCGCTC<br>TTGCGGGAGATATTGGATATTTTCTGGGGAGCCAAACCTCGTTGGA<br>AATTTTCAAAATTACATTGCCTGGCTTAGGGTTTGCTCTGCCATTAA<br>GCGTGGCAGTGAACGGCCTGACAATGGTGTTAGCAACGCTTGTGT<br>CATTTGTCATTGTCATGATCACACAATATTCCGTTGAGTACCTTGAC<br>GGTGACCCCCACCAAGCTCGTTTCTTTCGGTTATTAGCGTTCACCG<br>GCGGTTTCTTCTTACTCGTGGTAGTCAGCGGCAATATTGGGCTTTT<br>CACACTTGGCATCATTGCCACCGGCTTTTCACTGAATAAATTATTAA<br>AATTTTATAATACTCACCCTAAAGCCATAATGGCCGCGCATAAGAAG<br>AGCATCTTCACCAGAACGGCCGACTTATTTCTGGTTGGTGCTAGTG<br>CCTTGATAGGCTCGCATGTTGGTTCTCTGCAGTTTGACGCATTACG<br>TCAATTCGTGGCTCACGCAACCGAAATTCCTTTCGCCCTGCAAATA<br>GCTGCTTGGCTTATTATAGGCGCAGTGATTTTAAAGAGCGCACATT<br>TCCCGTTTCAGGGTTGGTTAATCCAAGTCATGGAAGCGCCGACCC<br>CCGTAAGTGCTCTCATGCACGCAGGGGTGGTGTACAGTGGCGCAA<br>TCATCGCTTTAAGAACTACGTCCTTACTGGTTCGCGTATCCGACGC<br>ACTCCTCTTCTTAGGCATCATGGGCCTTGCAACAGTGGTAATTGCC<br>AGTCTGGCGATGACGACTCAAACCGCCGTTAAATCTATGCTCGCAT<br>GGAGCACCACCGCACAACTCGGATTTATGTCCCTTGAGCTGGGTTT<br>AGGTTTATTTCCACTGGCCCTGTTGCATCTCATCGGGCATAGCTTA<br>TATAAAGCTCATGCGTTCTTAAGCTCTGGGTCCGTTCCGGATCAAT<br>TACGTCAGGTTCCCCCGGGTGGCAAGAAGATACCGAGCATGACCT<br>CATGGGTGATGGCTGTAGTGTTAGGCTTAATTATTTCCGGCGGCGG<br>CGCTTGGGTATTTGGTGACGACCCTCTGAAGGACCCCACATGGAT<br>GGCGTTGGTTGTAATTATAGCGGTCGCAGTGTCTCAGATTCTGATT<br>AAGGGATTCCAGTTTGACGCTATCATTGATCGCTTCGTCGCCTTCT<br>TAGTGGCCGTCTCAATGGGCTTCACTTACTTGATTTTGCATACCCT<br>GTTCGTTTGGGGATTTACGCGCGACATGACAGAAGCCATGATGCC<br>GATTACGACACCGTACCTTTGGCTGTTGGGCTTGAGCATGGCTAGT<br>TTCTTAATTCTGAGCTGGTTACAAGGACCAGGCCGAGCGCTGTTAC<br>CCCAGAAGATACAGCTGGTCTTGTTTACTCACCTGTATAATGGTCTT<br>TACGTAGACGTTTGGGTTGAACGCATCTCCCATCGATTTTGGAGCG<br>AGCGTGTCGGTGTGGAGTTACCACGGAAGAACTTGGCCGTCGAGT<br>CACTGCAGTCCATTGCCCGAAACCAAGACGCATTTGGTCAATCTGG<br>AGGTGTACAATGAGCCTCTGGATCGAC |
| *Acidithio-bacillus* | 2433 | ACTGACGGTCTCATGAGCTCTAACAAGACGGGAAACACAGAACGG<br>TTGTTTGATCGCGTCATTCGACGTATTGCCCCGGTGTGGCCATTAG |

| | |
|---|---|
| *caldus* | ATTCCTTCGTCGCTGTGAACCCGTATTGGGGCTTCGCAGATCAAAC |
| DabA | ATTCGCAGAAGCTGCCGCACATCTCCAGGGTAGTGTTGGAGAACG |
| | TGTGATTATGGACAGACGGTGGTACGCAGACCTGTTAAAGAATGGT |
| | AAATTGATGATGGCAGATATCCTTCAAGCAAGTAGTGATCTGGGTG |
| | TGGACTTAGACGAATACGGATGGCGTGCATACCTTAGAGAGCCGT |
| | CCGAACTGCCCATGAGACTGCCCCGTATTCCTGAGCTGATGGATC |
| | GACCAGGTCACCCAGCCGTGTCTCAGTATGTTGTAGAACAAATTTC |
| | GCGTTTTCTGGCGGCCTATTATGACCGCGGTCAATCACTGTGGAGT |
| | TTTCCTAAAGACCCCGCCGTTGGTCTGTTCGGGAGCTGGAGACAAT |
| | ATACTCTGCTGGACCGTACTCTTCGGGCTATGGGGTTAAAACATCT |
| | GCGTTCAGAACTTTTGACGGTCAGTAGCAACGCCGCATCTGCGCG |
| | CCAGTGGGCGCTGGAAACCCTGGCAATAGCACAACCCGCGGAAGA |
| | GGATTATCTGTTAGTGCTCTTGAAGTCGATTGGCGGCTGGGCGTCA |
| | TGGTGTCGGTATCTTCTTTGGCAAGCTGAATTGCAAGGCTCTACCA |
| | ACGCGGATTTGTTAGACCTGTTAACTATCCGTATGGTATGGGAAGG |
| | TTTGTTACGCAAAACTGCGGATGCACGTGTCCTCGAAAGATGGAGA |
| | CGTCAGATACATGCGTGGACCATTGATGATAAAGGTGTTGATTTAG |
| | AGGACGCTAGACGCGGCGAGGTTCTGCTGAGAGCGAGTGAAATCG |
| | CATATCGACGTCAAGTGGCGGCCTCAATCCGGCGGCAGCCCGCTA |
| | AGAAGAACCGTGCCCCGTTAATCCAAGCGGCGTTCTGTATTGATGT |
| | GAGAAGCGAAGTTTTCCGAAGACACTTGGAGTCAGTCATGCCCGA |
| | ATTGGATACAATAGGTTTCGCAGGCTTCTTTGGTGTACTGCTCGAC |
| | TATCAACGTAACGGGGACTACGCACCACGAACACAGACGCCAGTA |
| | TTACTGAACCCACAAGTTCATGTTACGGAAGACGCTCCAGGTGATG |
| | CCATACAAAGACGGTTCCGCCGTCTTCGTCGCGCAGCGGAATGGA |
| | AACATTTTAAACTGTCAGCCGCGTCATGCTTCTCGTTTGTCGAGAC |
| | AGCCGGGATAACGTATGTGGGACGGCTTTTGGCCGATACAATGGG |
| | CTGGCACCGGCCTAGCGTACACCCCGATGAAGCTGGTTTGTCACC |
| | GGAAGAACGTGCAAATCTTCGCTGTGTGCTGCCTAAGAGTTTGACG |
| | CTGGAACAAAGAATCGGTATGGCTGAATTTATCTTGACAGGTCTGG |
| | GATTAAATCGTGGGGTTGCACCAATCGTATTACTGGCGGGTCATGG |
| | CAGTAGCAATACTAACAACCCACACCGTGCGGGATTGGACTGTGG |
| | AGCTTGCGCCGGACAAACTGGGGAAGTTAATGCGAAAGCAGCCGC |
| | GGATTTATTAAATGACCCGGCAGTCCGCAAGGGTCTGGTCGAAAA |
| | GGGTTGGAACATTGACCCGCGTTGTTGTTTCTTGCCGGCATTACAT |
| | GATACAACTACTGACCGCGTAGAAATCTTAGGCGGCTTGGACAATC |
| | CAAAACTGGATACTGGCTTAGTGGCGGAACTCCAAGCCGCCCTTG |
| | ACAAGGCTGCAAATCTTACTCGCTTGGAGCGTATGCTGAGACTGGA |
| | ACCAGATCTCCGGGACCCAGAAACTGTGGCCCGTAATATGGAACA |
| | ACGTGGACGTGATTGGAGTCAGGTGCGCCCGGAGTGGGCCCTGG |
| | CGGGAAACGCAGCCTTTATAGCTGCGCCACGTTGGCGTACACGTG |
| | AAATGAACCTGGCCGGCCGTGCGTTTCTCCATGATTATGATATAGA |
| | AAAGGACCCCGACTTTGGGGTCCTTACATTAATTATGACAGCCCCA |
| | TTGGTCGTTGCTAATTGGATAAATCTGCAATACTATGGATCAATTGT |
| | AGATAATCAGAGACAGGGGTGTGGAAATAAGGTGTTACATAATGTA |
| | GTAGGTGGTACAATTGGTGTCTTGGAAGGCAACGGTGGTGACTTG |
| | CGCATCGGTTTAAGCGAGCAGTCACTGCGTGATTCTAACTCAGAGT |
| | TGCAACATGAGCCGCTTCGTCTGACAGCGTTTATCGAAGCTCCTAC |
| | GGATGCGATGGACCGTATAATAGCCAATAATGATGCCTTAGGGAGA |
| | CTGGTGAATAATCGCTGGTTATCAGTAGTTCAAATAGCTGCGGACG |
| | GTAGTCTTCGCGAACGTCGCTCCGAAGGACGGTGGGTATCAATTT |
| | GAAGCCTCTGGTCAGAT |

**Table S6.** Curated List of Diverse Uniprot DAB1 Sequences used for Conservation Analysis.

| DabA Entry | Gene Names | Organism | Associated DabB Entry | DabC present? |
|---|---|---|---|---|
| A0A059ZT80 | dabA Acaty_c0924 | *Acidithiobacillus caldus* (strain ATCC 51756 / DSM 8584 / KU) | A0A059ZTL4 | yes |
| A0A060UT06 | dabA AFERRI_20539 AFERRI_370022 | *Acidithiobacillus ferrivorans* | A0A060UN95 | yes |
| A0A0K6HSN6 | dabA Ga0061069_101421 | *Thiomonas bhubaneswarensis* | A0A0K6HSR8 | yes |
| A0A0P9C3F5 | dabA SAMN05661077_0175 | *Thiohalorhabdus denitrificans* | A0A1G5AAE3 | yes |
| A0A179BQF4 | dabA A4H96_00055 | *Acidithiobacillus ferrooxidans (Thiobacillus ferrooxidans)* | A0A179BQN6 | yes |
| A0A191ZGR6 | dabA A9404_06355 | *Halothiobacillus diazotrophicus* | A0A191ZGS1 | yes |
| A0ACD5HCP 7 | DUF2309 domain-containing protein, HHS34_009510 | *Acidithiobacillus montserratensis* (strain GG1-14) | A0ACD5HD5 5 | yes |
| A0A1D8IK76 | dabA BI364_01595 | *Acidihalobacter yilgarnensis* | A0A1D8IK93 | yes |
| A0A1D8KAU1 | dabA BJI67_14225 | *Acidihalobacter aeolianus* | A0A1D8KAS2 | yes |
| A0A1H3ZAL0 | dabA SAMN05660964_0111 2 | *Thiothrix caldifontis* | A0A1H3ZAN1 | yes |
| A0A1Z4VPS3 | dabA FOKN1_1083 | *Thiohalobacter thiocyanaticus* | A0A1Z4VPR4 | yes |
| A0AAE3CKJ7 | dabA HFQ13_12175 | *Igneacidithiobacillus copahuensis* (strain VAN18-1) | A0AAE2YRL6 | yes |
| A0A285P3L8 | dabA SAMN06265353_1106 | *Hydrogenobacter hydrogenophilus* | A0A285NYE1 | yes--fused |
| A0A2I1DKH3 | dabA B1757_09710 | *Acidithiobacillus marinus* | A0A2I1DKG2 | yes |
| A0A368HBX8 | dabA C4900_08585 | *Acidiferrobacter thiooxydans* | A0A1C2FZQ8 | yes |
| A0A401JHY3 | dabA SFMTTN_3545 | *Sulfuriferula multivorans* | A0A401JI09 | yes |
| A0A4R8IU21 | dabA EDC23_1865 | *Thiohalophilus thiocyanatoxydans* | A0A4R8IK62 | yes |
| A0A543Q3U1 | dabA DLNHIDIE_00855 | *Acidithiobacillus thiooxidans* ATCC 19377 | A0A543Q3T7 | yes |
| A0A656HG06 | dabA Thini_1858 | *Thiothrix nivea* (strain ATCC 35100 / DSM 5205 / JP2) | A0A656HC39 | yes |
| A0A6I6D7R3 | dabA GM160_03865 | *Guyparkeria halophila* | A0A6I6CXL2 | yes |
| A0A7C9P570 | dabA GZ085_05225 | *Sulfuriferula multivorans* | A0A7C9K1R7 | yes |

| | | | | |
|---|---|---|---|---|
| A0A7T4WF07 | dabA H2515_03250 | *Acidithiobacillus ferrivorans* | A0A1B9BXJ4 | yes |
| A0A845UBV2 | dabA GL267_05340 | *Acidithiobacillus ferrianus* | A0A845U389 | yes |
| A0A8I1MUZ6 | dabA J0I24_06505 | *Thiomonas arsenitoxydans* (strain DSM 22701 / CIP 110005 / 3As) | A0A8I1STW5 | yes |
| A0ABS5ZQJ3 | dabA HJG40_08035 | *Acidithiobacillus concretivorus* | A0ABS5ZRF7 | yes |
| A0ABS5ZVG3 | dabA HAP95_02765 | *Acidithiobacillus sulfurivorans* | A0ABS5ZVA8 | yes |
| A0ABU6FR22 | dabA OW717_10680 | *Acidithiobacillus ferriphilus* | A0ABU6FRH9 | yes |
| A0ABV0EK15 | dabA V6E02_11075 | *Thiobacter aerophilum* | A0ABV0EK25 | yes |
| A0ABV4TZC3 | dabA ACERLL_16645 | *Thiohalorhabdus methylotrophus* | A0ABV4TYQ5 | yes |
| A0A512LCI1 | dabA, TPL01_32400 | *Thiobacillus plumbiphila* | A0A512LC84 | yes |
| A0ABZ0YYS3 | dabA SR882_04640 | *Guyparkeria halophila* | A0ABZ0YYU6 | yes |
| A0ABZ1CMB8 | dabA VA613_13940 | *Thiobacillus sedimenti* | A0ABZ1CI67 | yes |
| A4BLJ5 | dabA NB231_15223 | *Nitrococcus mobilis* Nb-231 | A4BLJ3 | yes |
| C4FIQ8 | dabA SULYE_0448 | *Sulfurihydrogenibium yellowstonense SS-5* | C4FIQ7 | yes-fused |
| D0KZ77 | dabA1 Hneap_0907 | *Halothiobacillus neapolitanus* (strain ATCC 23641 / DSM 15147 / CIP 104769 / NCIMB 8539 / c2) *(Thiobacillus neapolitanus)* | D0KZ79 | yes |
| D3DIA8 | dabA HTH_1102 | *Hydrogenobacter thermophilus (strain DSM 6534 / IAM 12695 / TK-6)* | D3DIA7 | yes, fused |
| D3SP01 | dabA Thal_0253 | *Thermocrinis albus (strain DSM 14484 / JCM 11386 / HI 11/12)* | D3SP02 | yes, fused |
| D5X328 | dabA Tint_0113 | *Thiomonas intermedia (strain K12) (Thiobacillus intermedius)* | D5X326 | yes |
| D6CQJ7 | dabA THI_0132 THICB1_110318 | *Thiomonas arsenitoxydans (strain DSM 22701 / CIP 110005 / 3As)* | D6CQJ5 | yes |
| G0JL81 | dabA Acife_3126 | *Acidithiobacillus ferrivorans SS3* | G0JL83 | yes |

| | | | | |
|---|---|---|---|---|
| H5SBX7 | dabA HGMM_F07F09C38 | *uncultured Aquificia bacterium* | H5SBX8 | yes--fused |
| O67026 | dabA aq_863 | *Aquifex aeolicus (strain VF5)* | O67027 | yes--fused |
| Q0AHW4 | dabA Neut_0801 | *Nitrosomonas eutropha (strain DSM 101675 / C91 / Nm57)* | Q0AHW6 | yes |
| A0A1A6C0K6 | dabA Thpro_022334 | *Thiobacillus prosperus (strain DSM 5130/JVM 30709)* | A0A1A6C0J5 | yes |
| Q1QFS7 | dabA1 Nham_4323 | *Nitrobacter hamburgensis (strain DSM 10229 / NCIMB 13809 / X14)* | Q1QFS5 | yes |
| Q3SFK3 | dabA Tbd_2653 | *Thiobacillus denitrificans (strain ATCC 25259 / T1)* | Q3SFK1 | yes |
| A0A238D7K2 | dabA THIARS_70938 | *Thiobacillus delicatus (strain DSM 5494)* | A0A238D7N1 | yes |
| Q3SR42 | dabA1 Nwi_1990 | *Nitrobacter winogradskyi (strain ATCC 25391 / DSM 10237 / CIP 104748 / NCIMB 11846 / Nb-255)* | Q3SR40 | yes |
| W0DCR8 | dabA THERU_06545 | *Thermocrinis ruber* | W0DH74 | yes--fused |
| W9V6Y6 | dabA D779_1733 | *Imhoffiella purpurea* | W9VGL4 | yes |

**Movie S1 (separate file).** Remodeling of the lid region enclosing the active site (residues 474-509) and coordinated movement of DabB (residues 333-346).
